# Environmental DNA metabarcoding of invertebrate-incubated water supports WFD-compliant and animal-friendly bioassessment with added trait insights

**DOI:** 10.1101/2025.08.12.669652

**Authors:** Mandy Sander, Arne J. Beermann, Denis Brömmling, Dominik Buchner, Martina Weiss, Florian Leese

## Abstract

Environmental (eDNA) metabarcoding is recently being considered for bioassessment under the Water Framework Directive (WFD). While tissue-based (“bulk”) metabarcoding of macroinvertebrates produces comparable ecological status class values to morphology-based methods, eDNA-based metabarcoding often includes DNA signals from upstream sites, limiting its site-specificity. A promising alternative involves the incubation of locally collected invertebrates in water, followed by eDNA metabarcoding of the incubated water. In this study, we tested the suitability of two different approaches of the incubation strategy for ecological status assessment under the WFD. The first approach uses conventional multi-habitat sampling (MH) for the collection of the local macroinvertebrates and the second natural substrate exposures (NSEs) that are actively colonised. Using metabarcoding of the incubated water from both approaches (MH/NSE eDNA), we compared community composition, trait diversity and the derived ecological status classes (ESCs) to the local signal reference (MH/NSE bulk DNA) and the regional signal reference (stream water eDNA). For both approaches, community and trait composition were highly congruent between the incubated water and bulk samples. Trait composition did not differ between MH and NSE samples or among NSE types (wood-leaf vs. gravel). However, we detected small-scale spatial differences in the trait composition between NSEs placed at different flow regimes (pool vs. riffle). ESCs derived from all approaches (MH/NSE incubated water eDNA, MH/NSE bulk DNA and stream water eDNA) were highly similar and consistent with those from morpho-taxonomic assessments. Our findings support the incubation strategy using the conventional MH sampling as the most suitable approach for a minimally invasive, site-specific stream bioassessment within the context of the WFD.

## 1. Introduction

Ecological status assessment of freshwaters is routinely performed under the EU Water Framework Directive (WFD 2000/60/EC) using standardized sampling and analysis procedures. While the procedures can differ between countries, many aspects (e.g., assessments based on taxonomic composition and abundance, results reported as a ratio to a reference condition and the goal to achieve good ecological status) are similar, and the approaches are intercalibrated at European level (Birk et al. 2013). One of the key groups for assessing the ecological status of freshwaters are macroinvertebrates, which serve as biological quality elements (BQE) due to their sensitivity to stressors (Hering et al. 2003). The implemented traditional method for assessing the ecological state using freshwater macroinvertebrates as BQEs relies on morphological identification and abundance estimation of sampled macroinvertebrates. Then the composition of the found community is compared against an expected reference community for the water body type studied (Hering et al. 2004). Given the risk of wrong determination results and the time requirements for determination (Haase et al. 2004, 2010), many countries perform determination only to a higher taxonomic level, often genus or family.

A complementary or alternative approach to the traditional approach suggested for the implementation into the WFD is DNA metabarcoding of collected organismal bulk samples or environmental DNA (eDNA) (Leese et al. 2017; Hering et al. 2018; Brantschen et al. 2022; Macher et al. 2025). On the basis of eDNA traces shed from organisms into the water, comprehensive taxa lists can be derived using high-throughput sequencing and the subsequent comparison of the sequences to a reference database (Taberlet et al. 2012a, b; Valentini et al. 2016; Leese, Sander et al. 2021). The non-invasive easy and fast sampling, the potential of holistic assessments across different taxonomic groups (e.g., invertebrates, fish, diatoms) from one water sample and the high taxonomic resolution are some of the arguments supporting the implementation of eDNA metabarcoding into the WFD (Deiner et al. 2016; Stat et al. 2017; Macher et al. 2021a; Altermatt et al. 2025). For fish, ecological status classes derived from eDNA metabarcoding have been shown to be comparable to traditional methods (e.g., Hänfling et al. 2016; Pont et al. 2018; Pont et al. 2021). Thus, for fish, eDNA metabarcoding is currently being discussed for the implementation into regulatory assessment programs with the potential to replace harmful methods such as gillnetting, trawling or electrofishing (Hänfling et al. 2016; Hering et al. 2018; Pont et al. 2018). However, in contrast to metabarcoding of organismal bulk samples from locally collected macroinvertebrates (Elbrecht et al. 2017; Kuntke et al. 2020; Macher et al. 2025), implementing eDNA metabarcoding of macroinvertebrates into existing WFD regulatory biomonitoring poses several challenges (Hering et al. 2018; Altermatt et al. 2025). These challenges can be grouped into methodological, conceptual, perceptional and legal aspects (Leese et al. 2018). While many of the methodological, perceptional and legal aspects have been addressed in several of the published studies, a key aspect remains challenging: the lack of a clear site-specificity of the sample taken.

When trying to assess the local macroinvertebrate community with eDNA, transportation cannot be excluded, even over long distances exceeding 12 km (Deiner and Altermatt 2014). The retention and resuspension of DNA from the sediment can also impact the detection of the present macroinvertebrate community potentially resulting in the detection of false positives for local communities (Shogren et al. 2017). In contrast, traditional WFD protocols rely on locally collected samples, typically multi-habitat samples. This leads to discrepancies between taxa lists generated using eDNA metabarcoding and those obtained from local assessments based on physically collected animals, which can translate into deviant ecological status class assessments (Macher et al. 2018; Hajibabaei et al. 2019; Gleason et al. 2021; Pereira-da-Conceicoa et al. 2021; Múrria et al. 2024).

Evaluating local communities of a stream reliably, especially after a recent restoration or a local pollution event, is crucial for effective local impact assessments. While eDNA metabarcoding has proven useful for general ecological assessments (Li et al. 2018; Brantschen et al. 2021; Ji et al. 2022), its ability to accurately reflect the local community composition remains limited. To address this limitation, eRNA metabarcoding has been proposed as a more site-specific alternative (Cristescu 2019; Yates et al. 2021). However, recent studies show a low level of overlap and a lower species richness when comparing locally collected organisms with eRNA metabarcoding (Macher et al. 2024; Sander et al. in press). As a significant improvement for localized assessment, the results of Sander et al. (in press) suggest that incubating locally collected animals in water and using ‘eDNA enriched’ water for eDNA metabarcoding has a greater effect on the detection of the local community than the chosen molecule type (eDNA or eRNA) used for metabarcoding. The proposed strategy includes collecting local macroinvertebrate communities using multi-habitat (MH) kicknet sampling, as outlined by the WFD (Meier et al. 2006), incubating them in water, and subsequently analysing the incubated water (Sander et al. 2025). The prior incubation of the locally collected animals increased the overlap of species lists obtained by the eDNA enriched water from the incubation medium and the DNA extracted from the bulk sample of the locally collected animals by 20% compared to eDNA metabarcoding of a direct stream water sample (Sander et al. 2025; Sander et al. in press). Thus, the incubation strategy offers a more animal-friendly option by allowing the release of the animals after incubation, rather than relying on morphological identification or DNA analysis through metabarcoding of organismal bulk samples.

However, the use of kicknet sampling still involves an invasive method as it causes disruption to the streambed, which stresses and damages animals during the sampling process. Consequently, a second sampling approach for subsequent incubation of invertebrates was proposed, which uses precompiled substrates, so called natural substrate exposures (NSEs). The NSEs can be placed into the stream and are actively colonized by macroinvertebrates (Dumeier et al. 2018). Colonized NSEs can be carefully removed from the stream without disturbing the streambed. To account for spatial heterogeneity of the streambed, various types of NSEs can be introduced at multiple positions within a stream section. However, the NSEs may not replicate all microhabitats of a stream section with equal accuracy as some substrates such as sand or other elusive substrates cannot be replicated. Interestingly, when comparing both strategies, more species have been detected with the NSEs than with the MH sampling approach (Sander et al. in press). This is surprising since the NSE sampling approach offers a lower microhabitat and food source variety for macroinvertebrates than the broader range of natural stream habitats that are sampled using the MH sampling approach.

While the study by Sander et al. (in press) focused on the community composition and species richness detected by the two approaches of the incubation strategy, the suitability for ecological status assessment in the context of the WFD was not addressed. When applying the WFD in Germany, the ecological status of a stream is determined using the macroinvertebrate assessment approach “Perlodes” on the basis of taxonomic and abundance data, as well as traits associated with the specific taxon. Consequently, it is crucial for a sampling approach to be capable of reflecting the locally present trait diversity at a specific stream section to ensure the reliable assessment of the ecological status of a stream.

While eDNA metabarcoding of water samples has been shown to capture shifts in species trait composition for macroinvertebrates at small temporal scales (Sander et al. 2024), it has not been shown so far if the method can adequately detect trait differences at small spatial scales. In shallow and low-current transitional waters, eDNA metabarcoding can identify communities of species with different salinity preferences over small spatial scales (Pinna et al. 2024). However, most studies indicate that eDNA metabarcoding is not well-suited for assessing fine-scale spatial differences in taxonomic composition in streams (Deiner and Altermatt 2014; Gleason et al. 2021). Analysing eDNA enriched water after incubation of locally collected animals instead of water samples collected directly from the stream may improve the detection of small-scale spatial differences in small streams (Sander et al. in press).

We here applied the incubation strategy from Sander et al. (in press) to evaluate its potential value to support biomonitoring in the context of the WFD, specifically in relation to the German ecological status class assessment approach “Perlodes” that is based on macroinvertebrates. As a first step, we assessed the species trait composition of the invertebrate communities, which constitutes a key component for the general degradation module in “Perlodes”. For this, we first compared the trait diversity inferred from metabarcoding of incubated water eDNA samples to that inferred from multi-habitat (MH) and natural substrate exposure (NSE) bulk metabarcoding. Then, we determined whether the NSE approach provides a reliable estimation of macroinvertebrate trait composition in comparison to the conventional MH sampling approach. Furthermore, we examined potential differences in trait composition across NSE substrate types (wood-leaf vs. gravel) and habitat positions (pool vs. riffle) to determine the sensitivity of the NSE approach to habitat-specific differences. Finally, we evaluated whether the incubation strategy can reliably determine the ecological status of a stream.

Specifically, we aimed to answer the following questions:

1. Are species traits of invertebrate communities obtained by metabarcoding of incubated water eDNA samples and locally collected bulk samples from the NSEs comparable to those obtained by metabarcoding of incubated water eDNA samples and locally collected bulk samples from the MH approach?
2. Are ecological status class (ESC) values calculated using taxa lists from metabarcoding of incubated water eDNA samples and locally collected MH and NSE bulk samples comparable to the ESC values inferred from WFD assessments using classical morpho-taxonomic approaches for those streams?

In addition, we compared ESC values calculated using taxa lists from metabarcoding of stream water eDNA and eRNA to see how strongly different approaches diverge from each other and the classical morpho-taxonomy based assessment.

## 2. Material and methods

### 2.1 Sampling and experimental setup

We sampled four stream catchments each at two sites (eight sampling sites in total) located in Germany, North-Rhine Westphalia. The distance between the two sampling sites per catchment was at least 500 m. The two sampling sites of the Emscher catchment were located at two different tributaries of the Emscher, the Boye (BO) and the Borbecker Mühlenbach (BM) (Figure 1A). Before assembling the NSEs, the substrates for each sampling site were sampled on site to prevent contamination with pathogens or pesticides between streams and to provide natural substrates and food resources for the local macroinvertebrate community. Wood-leaf substrates consisted of 105 g wooden sticks cut to a length of 20 cm (diameter: 0.5 – 2 cm) tied together in three bundles and 20 g leaves (Figure A.1A). For each stream sticks and leaves were dried before weighing them and afterwards leaves were soaked in distilled water to prevent them from crumbling. Gravel substrates consisted of 2100 g gravel with a diameter of >1 cm to prevent the material from being flushed out of the nets (Figure A.1B). The substrates were placed in Nylon nets which measured 30 cm x 30 cm with a mesh size of 1 cm (Schutznetze24 GmbH, Aßlar-Berghausen, Germany). We prepared 16 wood-leaf NSEs and 16 gravel NSEs, resulting in 32 NSEs in total. The NSEs were placed in the stream four weeks before sampling to allow colonization from the local macroinvertebrate community. Considering the stream’s typology and based on the study of Dumeier et al. (2018), a colonization period of four weeks seemed to be optimal to increase species detection as longer colonization times may increase the risk of losing certain species due to the fast consumption of the food sources of the NSEs. At each sampling site, four NSEs were inserted, two wood-leaf and two gravel NSEs. One of each NSE type was placed at a section with a low current (pool) and the other at a section with a high current (riffle). Covering different flow regimes can improve the detection of taxa with diverging current preferences and hence also influence the occurrence of different feeding types (Cummins 2016). NSEs were secured to steel rods which were inserted 40 cm deep into the streambed (Figure 1B). To improve stability, the NSEs were further fixed using tent pegs. After two weeks, we checked the position and the fixation to the streambed for each NSE. The NSEs of the Emscher and Grüner Bach catchment were still in position and immersed in stream water, however, one NSE of the Boye was covered in sand, which had to be removed. All NSEs of the Ennepe and two NSEs of the Felderbach had to be relocated a few centimetres due to decreasing water levels.

**Figure 1.**
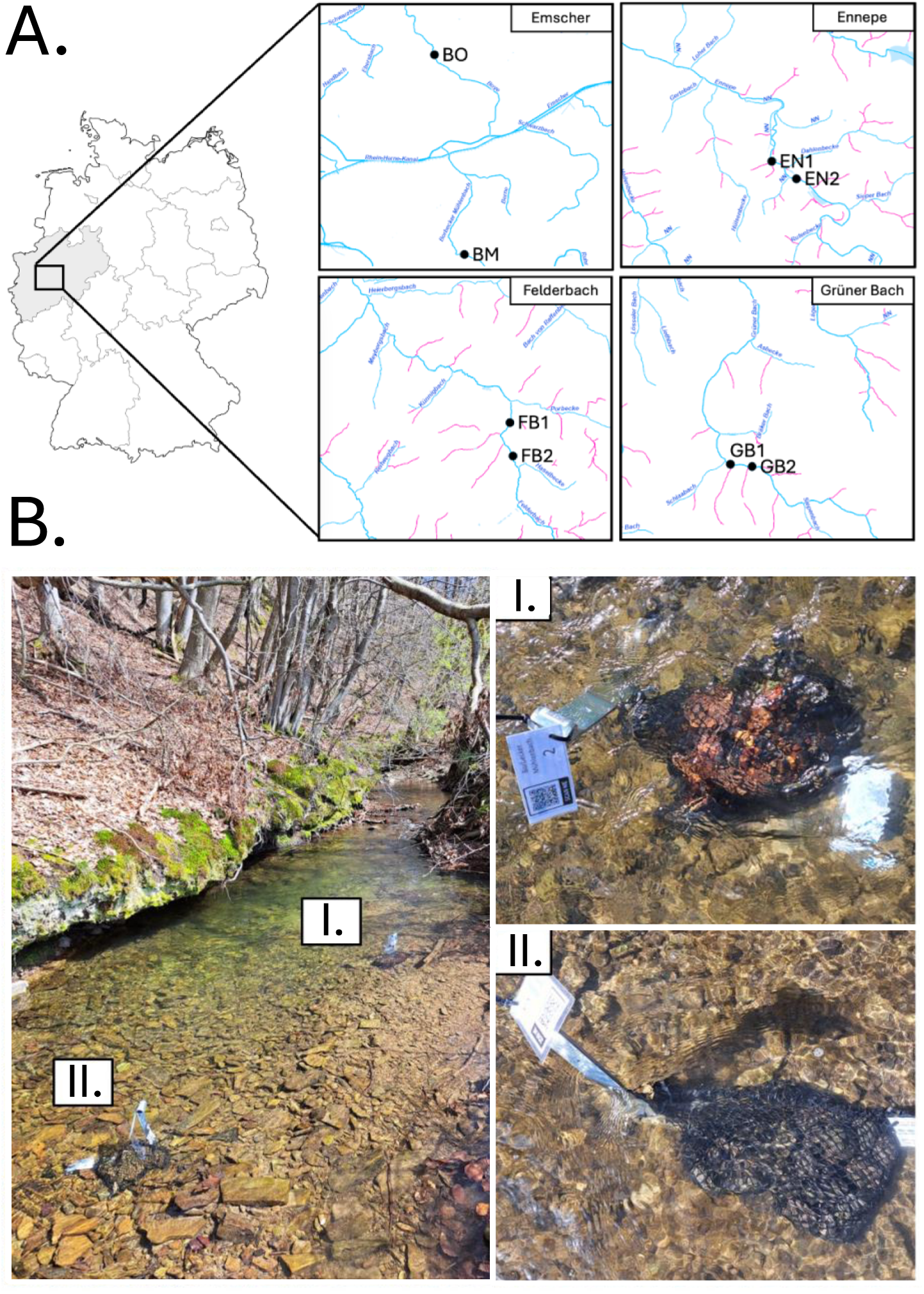
Sampling locations and overview of the NSE placement in the stream. A: Map of Germany showing the sampling locations of the four stream catchments Emscher (BO: 51°33’40”N; 6°55’59”E, BM: 51°26’16”N; 6°58’03”E), Ennepe (EN1: 51°16’59”N; 7°23’11”E, EN2: 51°16’49”N; 7°23’25”E), Felderbach (FB1: 51°21’11”N; 7°10’13”E, FB2: 51°20’55”N; 7°10’19”E) and Grüner Bach (GB1: 51°19’49”N; 7°40’51”E, GB2: 51°19’49”N; 7°41’02”E). Blue lines: surface waters; pink lines: additional surface waters without identification number. Altered after Sander et al. (in press). B: Placement of the natural substrate exposures (NSEs) in the stream illustrated at sampling site GB1 (left) and detailed picture of the two NSE types wood-leaf (I) and gravel (II) which were attached to a steel rod in the stream (right).

After another two weeks, sampling took place in April 2023. To minimize the impact from one sampling approach on the other, we started using the least invasive sampling approach first. At each sampling site we started with taking 500 mL water samples directly from the stream at the four sampling positions where the four NSEs were placed. Samples were taken downstream of the positions of the NSEs directed against the current to prevent interference with the NSEs. The stream water samples were directly filtered on site using a vacuum pump and 0.45 µm pore size cellulose nitrate membrane filters (diameter: 47 mm; Nalgene, New York, USA). After filtering, samples were stored on ice in the field. We refer to these samples as stream water samples.

Next, the NSEs were carefully removed with a net to avoid the loss of macroinvertebrates inhabiting the NSEs and to avoid the disruption of the streambed. Each NSE was transferred to a sterilised 20 L bucket filled with 10 L stream water from the respective site. After 15 min of incubation, the water in the bucket was stirred with a sterile stirring rod for 60 s to mix the water as well as the DNA in it. We let the suspended particles settle down for 4 min to reduce the risk of filter clogging. After a total of 20 min, we took a 500 mL water sample from the bucket which was filtered using the same vacuum pump and filter type as for the stream water samples. The filters were stored on ice and the animals from the bucket were preserved in 96% technical ethanol after removing larger sticks and gravel. We refer to these samples as NSE eDNA samples (water samples) and NSE bulk samples (organismal samples).

Lastly, we sampled a section of 20 m per sampling site using multi-habitat kicknet sampling according to the Water Framework Directive (WFD) (Meier et al. 2006) but with ten instead of 20 subsamples because of the reduced size and diversity of the streams (see supplementary Data S1 for information on the different habitat structures). Each stream section was divided into different microhabitats and the proportion of samples per habitat structure was noted down (Supplementary Data S1). Most streambeds were dominated by gravel except for the Boye streambed which was mainly covered in sand. Afterwards the whole kicknet sample was transferred to a sterilised bucket filled with stream water. Subsequent steps were identical to the ones used in the previous paragraph. We refer to these samples as MH eDNA samples (water samples) and MH bulk samples (organismal samples). For the MH eDNA sample of the Boye only 250 mL of the water from the incubation medium could be filtered due to filter clogging. This was likely due to the sediment that was flushed into the stream due to an ongoing construction next to the sampling site.

At each sampling site one air-contamination control sample was taken. To control for potential air-borne DNA we filtered air with a vacuum pump after all samples were filtered. All water samples and filters were immediately stored on ice in the field and later stored in a laboratory freezer at -80 °C. The animals were stored at room temperature in the laboratory and the preservative ethanol was exchanged with fresh 96% technical ethanol after 24 h. Figure A.2 gives a detailed overview of the number of samples taken per sampling approach and sample handling and processing.

### 2.2 Lysis and extraction

All laboratory steps were previously described in the study by Sander et al. (in press). Subsequently we shortly summarise the most important steps. For lysis of the bulk samples, we followed the protocol by Buchner et al. (2021a). Bulk samples containing the animals preserved in ethanol were homogenised with a kitchen blender (Mini Blender & Blender Smoothie, Homgeek, China). Per sample 1 mL of the homogenate was centrifuged to discard the supernatant ethanol. Afterwards the samples were ground at 2400 rpm for 2 min and lysed for 20 min at 65°C in a lysis medium of 900 µL TNES and 100 µL Proteinase K. The extraction of the bulk samples was done following the protocol from Buchner (2022a). Extraction was done in replicates and contained six extraction negative controls per 96-well plate. First, 500 µL binding buffer and 250 µL lysate per sample were mixed and 750 µL of the mixture transferred to the 96-well filter plate, which was placed on a vacuum manifold and filtered through the filter columns. The filters were then washed twice with 500 µL wash buffer. Then, the DNA bound to the filter column was eluted in 100 µL elution buffer. The extracted DNA was purified to avoid inhibition during PCR.

For the lysis of the DNA and RNA from the filters we followed the protocol from Buchner (2023). Filters were chopped into small pieces and each sample, including the 8 air-contamination control samples, was filled with 1000 µL of lysis buffer and ground at 2400 rpm for 2 min. For lysate clearing and pre-filtering 650 µL of the crude lysate was transferred to a pre-filter column and centrifuged for 10 min at maximum speed. Of the flow-through 600 µL were used for co-extracting DNA and RNA in replicates with 11 extraction negative controls per 96-well plate. Per replicate 300 µL were transferred to a filter column placed on a vacuum manifold. While the DNA was bound to the filter column, the RNA was kept as the flow-through of the filter in a new plate. RNA was precipitated by mixing the samples with 300 µL of 70% ethanol. For DNA extraction, the filter columns were washed, and the dried columns were subsequently eluted in 100 µL elution buffer. For the extraction of RNA, the filter columns were washed with RNA-specific wash buffers and eluted in 100 µL elution buffer. Afterwards the RNA eluate was digested with DNase using 9 µL of the extracted RNA mixed with 0.5 µL of 10 x ezDNase Buffer and 0.5 µL of ezDNase enzyme (ThermoFisher Scientific, Germany). The mixture was incubated for 2 min at 37°C. Afterwards, the RNA was transcribed to cDNA by adding 1 µL of Superscript IV VILO Master mix (ThermoFisher Scientific, Germany) and 9 µL of PCR-grade water to each sample and incubating the mixture for 10 min at 25°C followed by 10 min at 50°C and finally for 5 min at 85°C.

### 2.3 PCR and sample preparation

For metabarcoding the freshwater invertebrate primer pair fwhF2/fwhR2n targeting the COI gene region was used (Vamos et al. 2017). We used a two-step PCR approach with tagged versions of the fwhF2/fwhR2n primer pair with varying length and a universal tail in the first step (Leese, Sander et al. 2021) and primers matching the universal tail which included one of 96 i5/i7 indices and the P5/P7 Illumina adapter in the second step (Buchner et al. 2021a, b). The PCR reactions for the first step contained 1x Multiplex PCR Master Mix (Qiagen Multiplex PCR Plus Kit), 0.2 µM of each primer, 0.5x CoralLoad Dye and 1 µL purified DNA extract, which was filled up to a total of 10 µL with PCR-grade water. The cycling conditions consisted of the following steps: 95°C for 5 min, 20 cycles of 95°C for 30 s, 58°C for 90 s, 72°C for 30 s, with a final elongation at 68°C for 10 min. Subsequently, samples were purified to reduce potential primer-dimers that were formed during the first step (Buchner 2022b). The PCR reactions of the second PCR contained 1x Multiplex PCR Master Mix (Qiagen Multiplex PCR Plus Kit), 0.5 µM of each primer, 0.5x CoralLoad Dye and 2 µL of purified PCR product. The cycling conditions consisted of the following steps: 95°C for 5 min, 25 cycles of 95°C for 30 s, 61°C for 90 s, 72°C for 30 s, and a final elongation of 68°C for 10 min. Afterwards PCR products were normalized with magnetic beads (Buchner 2022c), pooled, purified and their DNA concentrations measured using a Fragment Analyzer™ Automated CE System (Advanced Analytical Technologies GmbH, Germany) with the NGS Standard Sensitivity Kit. To remove PCR bubbles from our library, which potentially formed after PCR, we performed a reconditioning PCR (Buchner 2024). The final library was sent to Macrogen Europe B.V. (Amsterdam, Netherlands) for sequencing on a single flow cell using an Illumina NovaSeq 6000 sequencer (150 bp paired-end reads, S4 flow cell) and 5% Phi-X spike-in.

### 2.4 Data analysis

For bioinformatic processing of the sequences we used the APSCALE pipeline version 1.6.3 (Buchner et al. 2022), which is based on VSEARCH and cutadapt (Martin 2011; Rognes et al. 2016). We used standard settings to process the demultiplexed sequences and clustered OTUs with a 97% similarity threshold. Filtered sequences were taxonomically assigned via BOLDigger version 2.1.1 to a sequence deposited in BOLD (Buchner and Leese 2020) with the following thresholds: ≥98% for species level, ≥95% for genus level, ≥90% for family level, ≥85% for order level, <85% for class level. These are the default settings in BOLDigger where the species threshold is based on the study of Hebert et al. (2003). We merged the extraction replicates per sample and retained all OTUs with reads in both replicates. To account for false positives and tag switching, the maximum number of reads among negative controls per OTU was subtracted from each OTU in the samples. We retained all OTUs with an assignment to at least family level for further analysis. The dataset was filtered for macroinvertebrates and all terrestrial taxa were removed after comparing our taxa list to the entries deposited in freshwaterecology.info (Schmidt-Kloiber and Hering 2015, 2024). Lastly, we used TaxonTableTools version 1.4.8 (Macher et al. 2021b) to filter the taxa lists to the taxa occurring in the operational taxa list (OTL) “Perlodes” used in Germany for Water Framework Directive (WFD) assessments.

### 2.5 Species trait data

For all detected taxa we used trait data from freshwaterecology.info (Schmidt-Kloiber and Hering 2015, 2024) if available to compare trait diversity between multi-habitat (MH) and natural substrate exposure (NSE) samples as well as between NSE types (wood-leaf vs. gravel) and between NSE position in the streams flow regime (pool vs. riffle). For this we used the original taxa list filtered to aquatic macroinvertebrates and not the OTL-filtered list to access a greater number of species-and genus-level data. We used species-level trait values where available and we used data from a higher taxonomic level, such as genus- or family-level trait values if species-level values were missing. For taxa with no trait values at the species-, genus-, or family-level, we used an averaging “gap-filling” procedure whereby values were averaged across species within the genus if species-level values were available and across all taxa within the family if not (Kunz et al. 2022). We assigned the traits feeding types and microhabitat preference for the comparisons between MH and NSE samples and between NSE types and used feeding type preference and current preference for the comparison between NSE riffle and pool samples. For the trait analysis we used both taxa lists from metabarcoding of the locally collected bulk samples and of eDNA enriched water by incubation of the locally sampled bulk. For statistical analysis we used relative trait composition per sampling site.

### 2.6 Ecological status class assessment

For the ecological status class assessment (ESC) we used the OTL-filtered taxa list generated by metabarcoding and formatted the list to be usable for ESC calculation via the online tool “Perlodes” (https://gewaesser-bewertung-berechnung.de/index.php/home.html). ESC values were calculated based on presence-absence data. For ESC calculations the stream type of the sampled stream section is an important factor. According to LAWA typology all sampled streams belonged to stream type 5 (coarse stony substrates of high silicate concentration) except for the Emscher tributaries. The Borbecker Mühlenbach belonged to stream type 6 at the sampled site (stream type 6: small fine substrate dominated calcareous highland rivers) and the Boye to stream type 14 (stream type 14: small sand-dominated lowland rivers) (LAWA-Typology, Dahm et al. 2014). To compare our data to morpho-taxonomic assessments, we extracted ESC values from ELWAS-WEB (https://www.elwasweb.nrw.de/elwas-web/index.xhtml, last accessed: 31.03.2025) which contains monitoring data on the water network, on the principles and results of water monitoring and on the ecological status according to the WFD. We used ESC values from the most recent sampling campaign, which was conducted between 2019 and 2021. We used either the closest monitoring site of the sampled stream within a radius of 1 km to our sampling sites or the only sampling site available within the sampled stream. The only exception was the Boye sampling site, where ESC values were only available from older sampling campaigns or from sites which were farther away. Therefore, for the Boye we used the ESC values calculated by a sampling campaign conducted by expert taxonomists at the site, which ranged between a “good” and “moderate” status. For the Grüner Bach we used the ESC values from the most recent sampling campaign (2019-2021) assigning a “high” status as well as from an older campaign, which assigned a “good” status, because there was no taxa list available to verify the ESC values of the newest sampling campaign. For the status class assessment of the NSEs, we calculated the ESC value based on the combined taxa list of the four NSEs per sampling site. We also calculated the ESC values for each NSE separately to see how much the results vary from the combined results. Furthermore, we conducted an intercalibration on our results from the two modules “Saprobic index” and “General Degradation” according to the method used by Macher et al. (2025). For stream type 6, no intercalibration could be conducted, as the stream type was not included in the intercalibration method used.

### 2.7 Statistical analysis

All statistical analyses and plots were generated with R version 4.0.5 (R Core Team 2021) and the ggplot2 package (Wickham 2011). We tested if beta-diversity values based on Jaccard dissimilarity differ between metabarcoding of the locally collected bulk samples (MH/NSE), the incubated water eDNA samples (MH/NSE) and the direct stream water eDNA/eRNA samples. For this, we calculated Jaccard dissimilarity using the betapart package (Baselga et al. 2018) and checked if the beta-diversity values of the samples were normally distributed using a manual inspection based on Q-Q plots and a Shapiro-Wilk test. Additionally, we tested the samples for homogeneity with a Levene test using the R-package car version 3.1-1 (Fox and Weisberg 2019). The samples were normally distributed and showed no heterogeneity in their variance. Therefore, an ANOVA (Analysis Of Variance) and a subsequent Tukey’s HSD test were performed to check which samples differed significantly from each other (see Supplementary Data S2 for details). Additionally, we compared only beta-diversity values between samples from a sampling site with a stream type 5 classification according to LAWA to test for the effect of stream type.

For the comparison of trait distributions between the two incubation sampling approaches MH and NSE and between NSE types and position within the stream flow regime, a PERMANOVA (Permutational Multivariate Analysis Of Variance) was performed using the vegan package (Oksanen et al. 2022). To test if differences between groups can be also attributed to differences in the sample’s variance, we performed a PERMDISP (Permutational Multivariate Analysis of Dispersion) using the vegan package, which is a multivariate extension of the Levene test. The PERMDISP did not support that the differences between groups were caused by the sample’s variance. We tested the effect of the sampling approach (MH vs. NSE)/NSE type (wood-leaf vs. gravel)/NSE stream habitat (pool vs. riffle) on the proportion of the different categories per trait using a constrained permutation design, where the permutation occurs on the repeated measures within each stream catchment (see Supplementary Data S2 for details). Subsequently, we tested which categories within each trait differed significantly between comparisons by performing a paired samples Wilcoxon Signed Rank test followed by a False Discovery Rate (FDR) p-value correction.

Lastly, we performed an indicator species analysis using presence-absence data using the function multipatt() in the package indicspecies (De Cáceres et al. 2010) to identify species that were associated to either pool or riffle habitats (see Supplementary Data S2 for details). We performed the analysis for each trait category separately to achieve an accurate weighting within the trait categories and consequently identify indicators of low-abundant categories. We chose the indicator value index (Dufrêne and Legendre 1997) rather than a correlation-based index, as the former is more suited for biomonitoring aspects (see Sander et al. 2024).

## 3. Results

### 3.1 General metabarcoding statistics

The metabarcoding analysis yielded 334,260,739 raw reads, which clustered into 23,381 OTUs after bioinformatic quality filtering (see supplementary Data S3 and Appendix A Table A.1 for more details on read distribution among sampling approaches). The 63 negative controls contained 659,594 reads, representing 0.22% of the total reads. Only one replicate of one of the extraction negative controls (i.e., 1/126) on the bulk metabarcoding extraction and PCR plate contained an exceptionally high read number with 570,292 reads. We subtracted reads found in all negative controls from our samples, including this negative control replicate, and thus do not expect an impact of this on the analysed samples. One replicate of one of the eRNA samples contained no reads after quality filtering, however, the other replicate contained sufficient reads for subsequent statistical analysis, and we decided to keep the sample in the analysis using only this replicate.

For ecological analysis, we retained 1498 OTUs assigned to 843 taxa and 682 unique species after filtering the taxonomic list to aquatic macroinvertebrates with ≥90% similarity (see Supplementary Data S4). Of the 843 taxa, 236 taxa had a record in the operational taxa list (OTL) used in Germany for Water Framework Directive (WFD) assessments. The highest number of OTUs (1117/1025), species (515/439) and OTL taxa (189/172) was detected by the NSE sampling approach (incubated water eDNA/bulk DNA respectively) followed by the stream water eDNA sampling approach, while the stream water eRNA sampling approach yielded the lowest richness. However, OTL taxa richness was comparable across sampling approaches and ranged between 123 and 189 taxa (Figure 2). Incubating the locally collected animals increased the similarity between the OTL-filtered lists obtained by metabarcoding of eDNA and bulk DNA from 55% to 69% for the MH sampling approach and from 60% to 73% for the NSE sampling approach (Figure 2).

**Figure 2.**
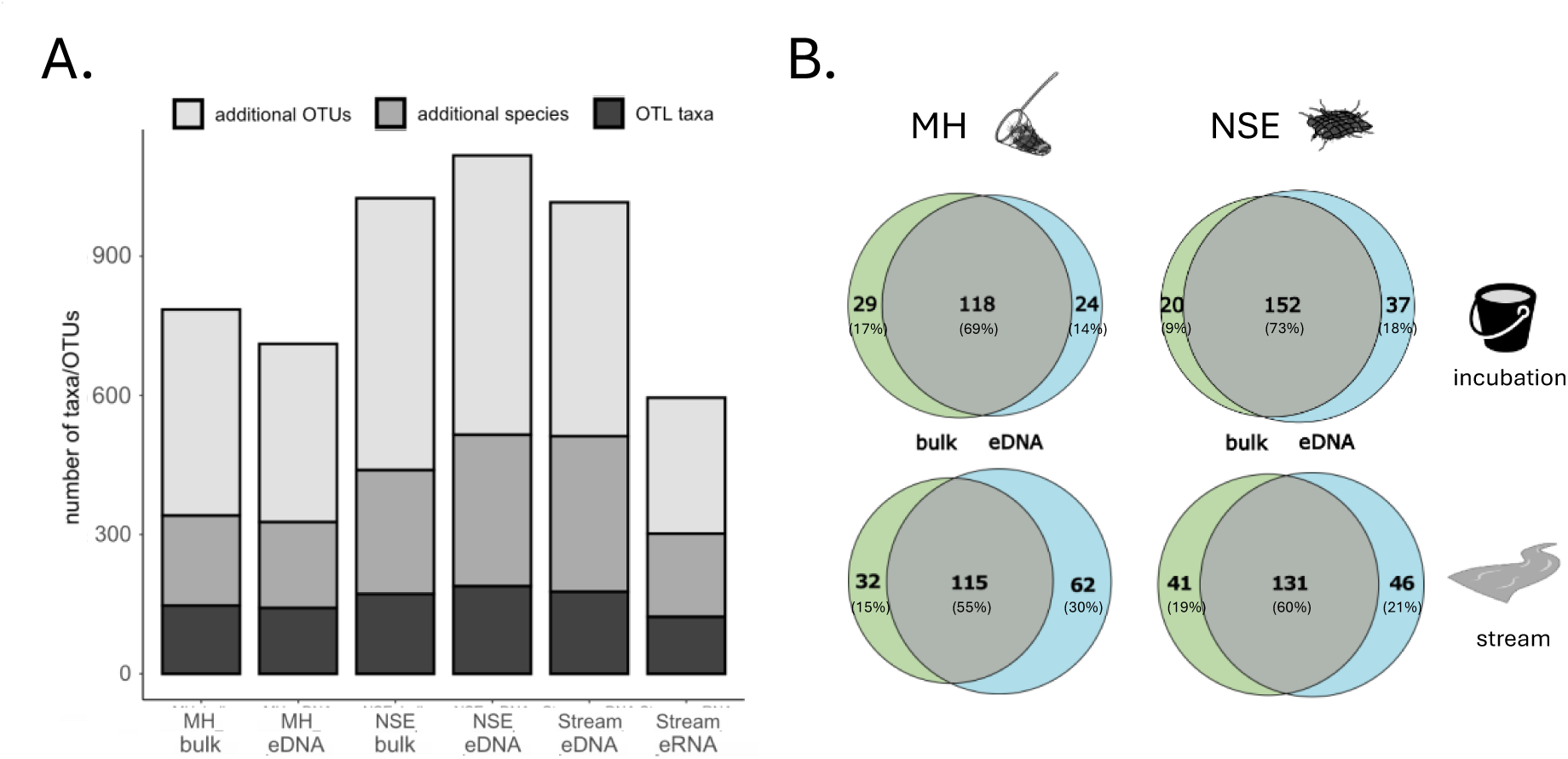
Comparison of the taxa/OTU richness of the different sampling approaches: multi-habitat sampling (MH bulk and eDNA), natural substrate exposures (NSE bulk and eDNA) and direct stream sample (eDNA and eRNA). A: number of OTL taxa, species and OTUs detected per sampling approach. Dark grey: taxa included in the operational taxa list (OTL), mid-grey: additional species detected by metabarcoding but not included in the OTL, light grey: additional OTUs detected by metabarcoding but not included in the OTL or assigned to one species. B: Venn diagrams showing the number of taxa from the OTL-filtered list shared between bulk DNA metabarcoding and incubated water eDNA metabarcoding (grey) and the exclusively detected taxa (green for bulk and blue for eDNA). Top: Incubation strategy, bottom: direct stream samples, left: comparison to the locally sampled bulk from the MH sampling approach, right: comparison to the locally sampled bulk from the NSE approach.

### 3.2 Comparison of species community composition

We compared the community composition using metabarcoding of the eDNA enriched water via the incubation strategy (MH and NSE) to the local signal reference (MH/NSE bulk DNA) and the regional signal reference (stream water eDNA). Across all comparisons, community composition inferred from the incubated water eDNA was most similar to the respective bulk sample (Figure 3). For MH bulk samples, this difference was not statistically significant, nor were the differences to the stream water (ANOVA: F-value = 2.356, p-value = 0.119, numerator degrees of freedom (ndf) = 2, denominator degrees of freedom (ddf) = 21). However, for the NSE sampling approach, Jaccard dissimilarity between stream water eDNA and bulk DNA was significantly higher than between incubated water eDNA and stream water eDNA or incubated water eDNA and bulk DNA (ANOVA: F-value = 8.756, p-value = 0.002, ndf = 2, ddf = 21). For samples collected at streams classified as type 5 according to LAWA typology, all comparisons were significantly different, except between the Jaccard dissimilarity values from the stream water comparisons (ANOVA: MH: F-value = 27.38, p-value = <0.001, ndf = 2, ddf = 15; NSE: F-value = 14.99, p-value = <0.001, ndf = 2, ddf = 15). Stream water eDNA samples were most dissimilar to the bulk DNA samples analysed (Figure 3). Stream water eRNA had the highest dissimilarity to the bulk DNA samples, while stream water eDNA and bulk DNA samples had the lowest dissimilarity, which was significant for the NSE samples of stream type 5 (ANOVA: MH: F-value = 1.977, p-value = 0.163, ndf = 2, ddf = 21; MH type 5: F-value = 2.451, p-value = 0.12, ndf = 2, ddf = 15; NSE: F-value = 2.55, p-value = 0.102, ndf = 2, ddf = 21; NSE type 5: 3.907, p-value = 0.043, ndf = 2, ddf = 15) (Figure A.3).

**Figure 3.**
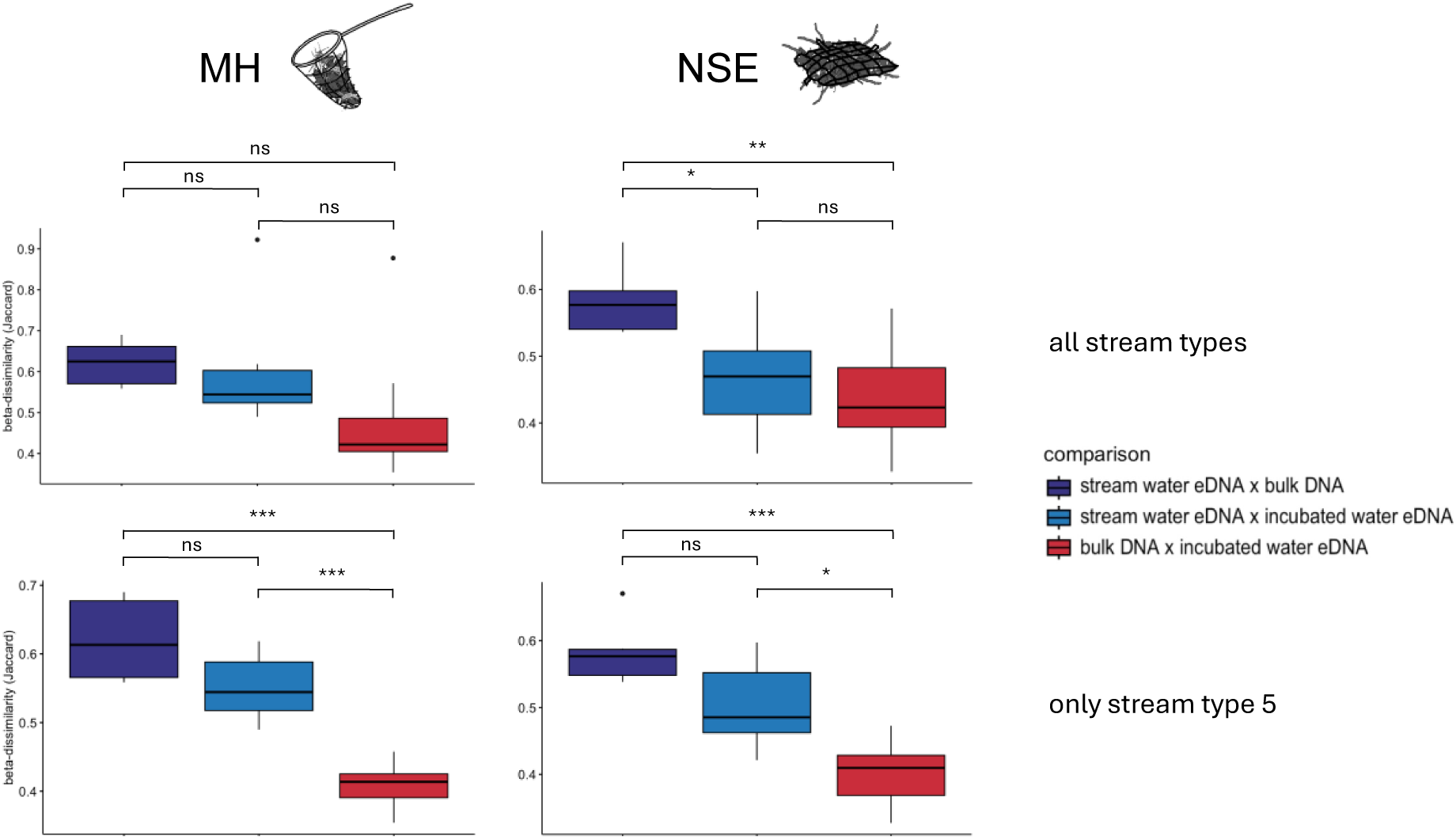
Comparison of species community composition based on Jaccard dissimilarity between incubated water eDNA, bulk DNA and stream water eDNA. Left: multi-habitat kicknet sampling approach (MH), right: natural substrate exposure sampling approach (NSE). Top: all streams, bottom: only streams assigned to stream type 5 according to LAWA typology. Significance between comparisons based on the results of the Tukey’s HSD test is presented as ns = not significant; * = p ≤ 0.05; ** = p ≤ 0.01; *** = p ≤ 0.001.

### 3.3 Comparison of species trait composition

We analysed the species traits microhabitat preference, feeding type and current preference and tested if trait composition differed between the two sampling approaches MH and NSE and between NSE types (wood-leaf and gravel) and NSE position in stream sections that differed in their habitat characteristics (riffle and pool).

For the species detected in the MH and NSE samples, a trait for microhabitat preference and a feeding type could be assigned to 541 (91%) and 557 (94%) of the 592 detected species, respectively (Supplementary Data S5). Most traits (microhabitat preference/feeding type) were assigned at species-level (364/288), followed by genus-level (132/153), family-level (47/97) and class-level (13 Ostracoda species assigned to a feeding type) with only 1 and 3 traits assigned through gap-filling, respectively (Supplementary Data S5). For current preference we only took the samples of the NSE approach into account. In total 538 (95%) of the 566 species detected with the NSE approach had a species trait assigned. Of these 232 were assigned to species-level, 150 to genus-level, 104 to family-level, 13 to class-level (all Ostracoda) and 9 were assigned using gap-filling (Supplementary Data S5).

We found no significant difference between the proportions of species traits for the MH and NSE sampling approach neither for microhabitat preference (incubated water eDNA PERMANOVA: F-value = 0.823, p-value = 0.094, ndf = 1, ddf = 14) nor feeding type (incubated water eDNA PERMANOVA: F-value = 0.426, p-value = 0.469, ndf = 1, ddf = 14; bulk DNA PERMANOVA: F-value = 0.542, p-value = 0.375, ndf = 1, ddf = 14). However, microhabitat preference significantly differed between bulk DNA samples of the MH and NSE approach, but none of the paired tests indicated a significant difference between single trait categories (bulk DNA PERMANOVA: F-value = 1.792, p-value = 0.008, ndf = 1, ddf = 14). Similarly, species trait composition did not differ between the two NSE types, wood-leaf and gravel (microhabitat preference: incubated water eDNA PERMANOVA: F-value = 0.573, p-value = 0.153, ndf = 1, ddf = 30; bulk DNA PERMANOVA: F-value = 1.213, p-value = 0.124, ndf = 1, ddf = 30; feeding type: incubated water eDNA PERMANOVA: F-value = 0.394, p-value = 0.64, ndf = 1, ddf = 30; bulk DNA PERMANOVA: F-value = 2.007, p-value = 0.127, ndf = 1, ddf = 30). Here, even after analysing the trait composition between NSE types for each stream separately, no differences were detected. In contrast, the position of the NSEs in the stream’s flow regime (riffle and pool) had a significant effect on the species trait composition of both, the current preference of species and their feeding type.

The analysis of incubated water eDNA and bulk DNA samples revealed significant differences in the composition of the species trait current preference of riffle and pool habitats. When each stream was analysed individually, a significant effect of stream habitats was observed for both sample types (incubated water eDNA PERMANOVA: F-value = 1.458, p-value = 0.002, ndf = 1, ddf = 30; bulk DNA PERMANOVA: F-value = 10.255, p-value = 0.001, ndf = 1, ddf = 30). However, the effect size of the pseudo-F statistic of 1.458 was small for the incubated water eDNA samples, suggesting a weak distinction between the different flow regimes. In contrast, the bulk DNA samples exhibited a more pronounced pattern of species distribution in relation to stream position.

A higher proportion of species preferring higher currents was found in samples taken at riffles (59%/53% compared to 51%/50% at pools for bulk DNA and incubated water eDNA respectively), while a higher proportion of species preferring lower currents was detected in the pool samples (27%/27% compared to 20%/23% at riffles for bulk DNA and incubated water eDNA respectively). This pattern was found for both bulk DNA and incubated water eDNA but was more pronounced in the former (Figure 4A and B). The indicator species analysis revealed that certain species were associated with specific flow regimes. Species preferring standing waters (limnophilic and limno- to rheophilic) were all associated with the pool samples and were planktonic species, such as Copepoda and Ostracoda (Table A.2). Similarly, the seven indicator species adapted to higher currents (rheophilic) were all associated with the riffle samples. The only exception was the parasite *Epoicocladius ephemerae* (Kieffer, 1924), which was associated with the pool samples (Table A.2). In contrast, no indicator species was assigned to the current preference categories on the extreme ends (limnobiont and rheobiont). However, the rheobiont genus *Rhyacophila* Pictet, 1834 was, apart from one species, exclusively found in the riffle samples, albeit at a low frequency among samples. Three of the *Rhyacophila* species were only detected in the incubated water eDNA samples, while one species was detected in the bulk DNA samples.

**Figure 4.**
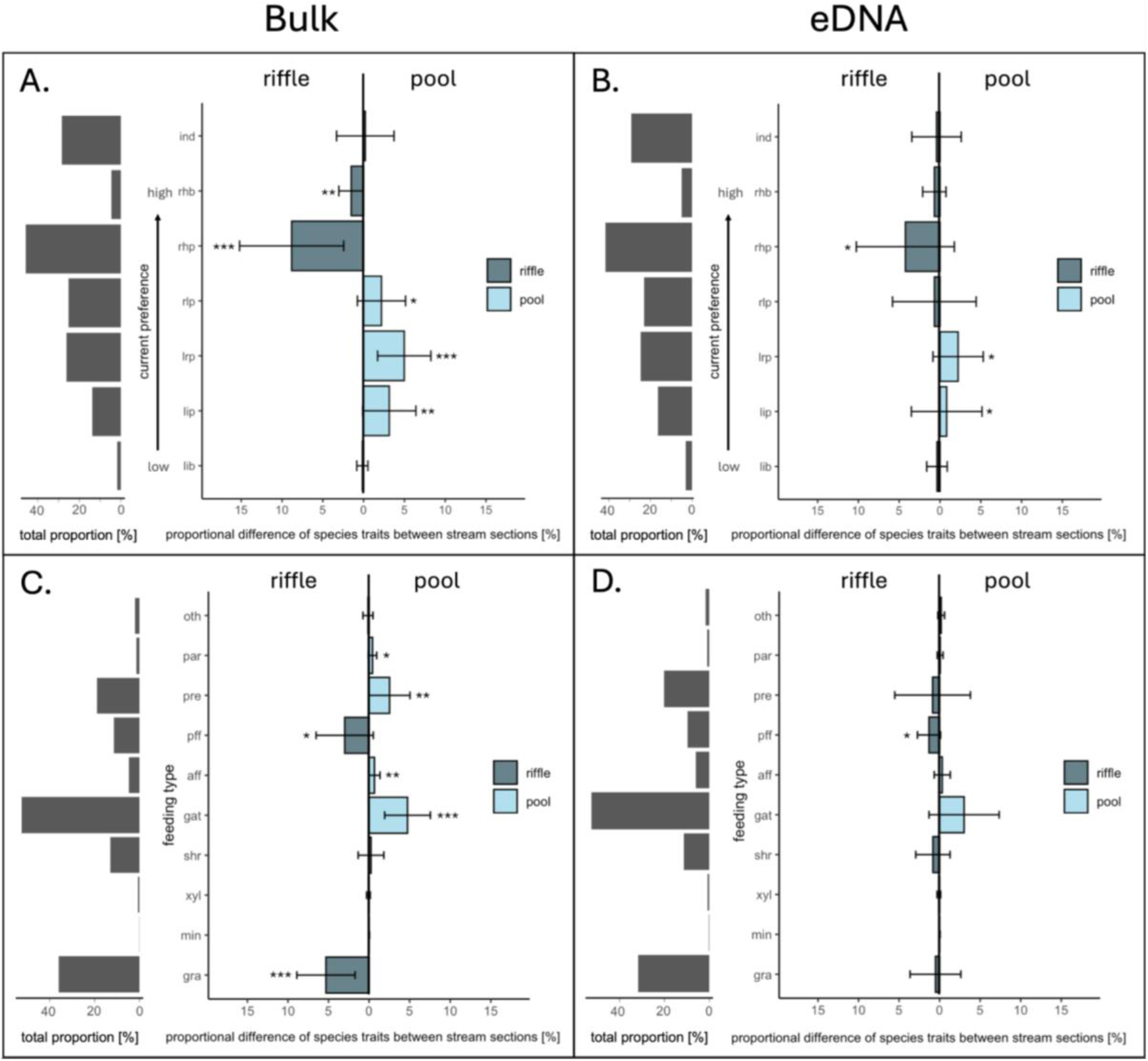
Stream current preference and functional feeding groups of macroinvertebrate species sampled in riffle and pool stream sections analysed with bulk DNA and incubated water eDNA metabarcoding. A: total proportion of each current preference category (left) and proportional difference between riffle and pool samples for each current preference category (right) for bulk DNA samples; B: total proportion of each current preference category (left) and proportional difference between riffle and pool samples for each current preference category (right) for incubated water eDNA samples; C: total proportion of each feeding type (left) and proportional difference between riffle and pool samples for each feeding type (right) for bulk DNA samples; D: total proportion of each feeding type (left) and proportional difference between riffle and pool samples for each feeding type (right) for incubated water eDNA samples. Dark blue: higher proportion in riffle samples, light blue: higher proportion in pool samples. Significance between comparisons calculated by pairwise Wilcoxon Signed Rank tests is presented as: * = p ≤ 0.05; ** = p ≤ 0.01; *** = p ≤ 0.001.

Feeding type composition differed significantly between riffle and pool stream habitats for the analysis of incubated water eDNA and bulk DNA. For both sample types a significant effect of stream habitat was observed when each stream was analysed individually (incubated water eDNA PERMANOVA: F-value = 2.2592, p-value = 0.021, ndf = 1, ddf = 30; bulk DNA PERMANOVA: F-value = 13.273, p-value = 0.001, ndf = 1, ddf = 30). In the incubated water eDNA and bulk DNA samples passive filter feeders were more prevalent in the riffle section samples (Figure 4C and D). In addition, grazers were also more common in the bulk DNA riffle section samples. In contrast, gatherer, active filter feeders, predators, and parasites were more frequently detected in the pool section samples (Figure 4C). Of the 15 indicator species identified for the feeding type analysis, 12 were also identified as indicator species for the current preference analysis and 5 species were identified as indicators for two feeding types (Table A.3). Among those, 10 were associated with the riffle section samples and 5 with the pool section samples. Grazer and passive filter feeder indicator species were predominantly found in the riffle samples, whereas the active filter feeder and the parasite *E. ephemerae* were associated with pool samples. Gatherer indicator species were equally represented in the riffle and pool section samples, and three out of four predator indicator species were associated with the riffle section samples, while only one was associated with the pool section samples (Table A.3).

### 3.4 Comparison of ecological status class values

We compared ecological status class (ESC) values calculated from taxa lists generated by metabarcoding to available ESCs based on morpho-taxonomic assessment. All streams, except the Emscher tributary Borbecker Mühlenbach (ESC value = poor), were assessed to be in a moderate or good ecological state, while the two Grüner Bach sampling sites were in a good to high ecological state according to ELWAS (Figure 5). ESCs calculated based on metabarcoding data were either identical to the morphology-based values from ELWAS or differed only by one class, except for the stream water eRNA samples, which had in general lower ESC values (Figure 5). In addition, neither the incubation of the locally collected bulk samples nor the sampling approach (MH or NSE) influenced the calculated ESC (Figure 5). Furthermore, intercalibration of the ESC values did not lead to deviations of more than one status class from the original values or to a generally higher agreement to the morpho-taxonomic assessments (Figure A.4).

**Figure 5.**
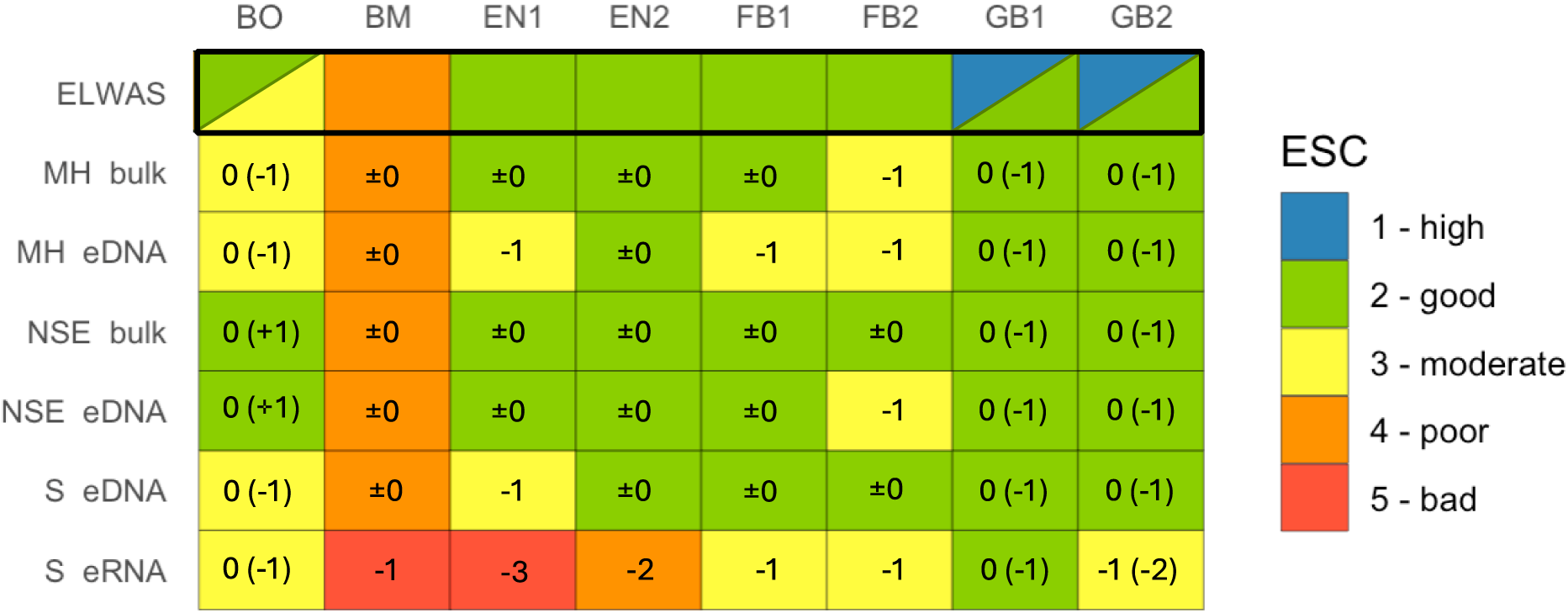
Ecological status class (ESC) values calculated from taxa lists generated via metabarcoding and morphology-based taxa lists. ESC values range from 1 = high to 5 = bad. Values in brackets indicate the divergence of the value calculated via metabarcoding from the value stored in ELWAS. For sampling sites BO no recent morphology-based value was available at ELWAS. The values displayed for the BO site are based on values extracted from ELWAS from the years 2012-2014. In addition, data from routine monitoring at this site reported an ESC value between 2 and 3. For the two sites GB1 and GB2, the most recent entry from ELWAS reports an ESC value of 1, however, no taxa list was available to validate the result. Therefore, we also included the older value stored with taxa lists in ELWAS, which report an ESC of 2.

Separate analysis of the four NSEs per sampling site revealed slight deviations among NSEs, with greater divergence from the morpho-taxonomic assessments than when taxa lists were combined (Figure A.5, 6). These discrepancies did not systematically differ based on NSE type or placement at different stream habitats. Particularly in the Emscher tributaries, deviations between NSEs were more pronounced (Figure A.5, 6).

Although ESC values calculated based on incubated water eDNA, bulk DNA and stream water eDNA samples were comparable, the detection of Ephemeroptera, Plecoptera and Trichoptera (EPT) species among sampling approaches was not congruent. For the order Ephemeroptera, this discrepancy in species detection was less pronounced among sampling approaches (Figure 6). However, many Plecoptera and especially Trichoptera species were exclusively detected using the two incubation strategies (MH and NSE). Here, at sampling sites FB2 and GB1 only one and two Trichoptera species, respectively, were detected in the stream water eDNA samples (Figure 6). The two incubation strategies shared most of the EPT species but overall, the NSE sampling approach detected more species than the MH sampling approach (Figure 6). Although the two sampling sites BO and BM of the Emscher catchment showed a significantly lower EPT species richness compared to the other streams (Figure 6), the Boye still had a moderate to good ESC value assigned.

**Figure 6.**
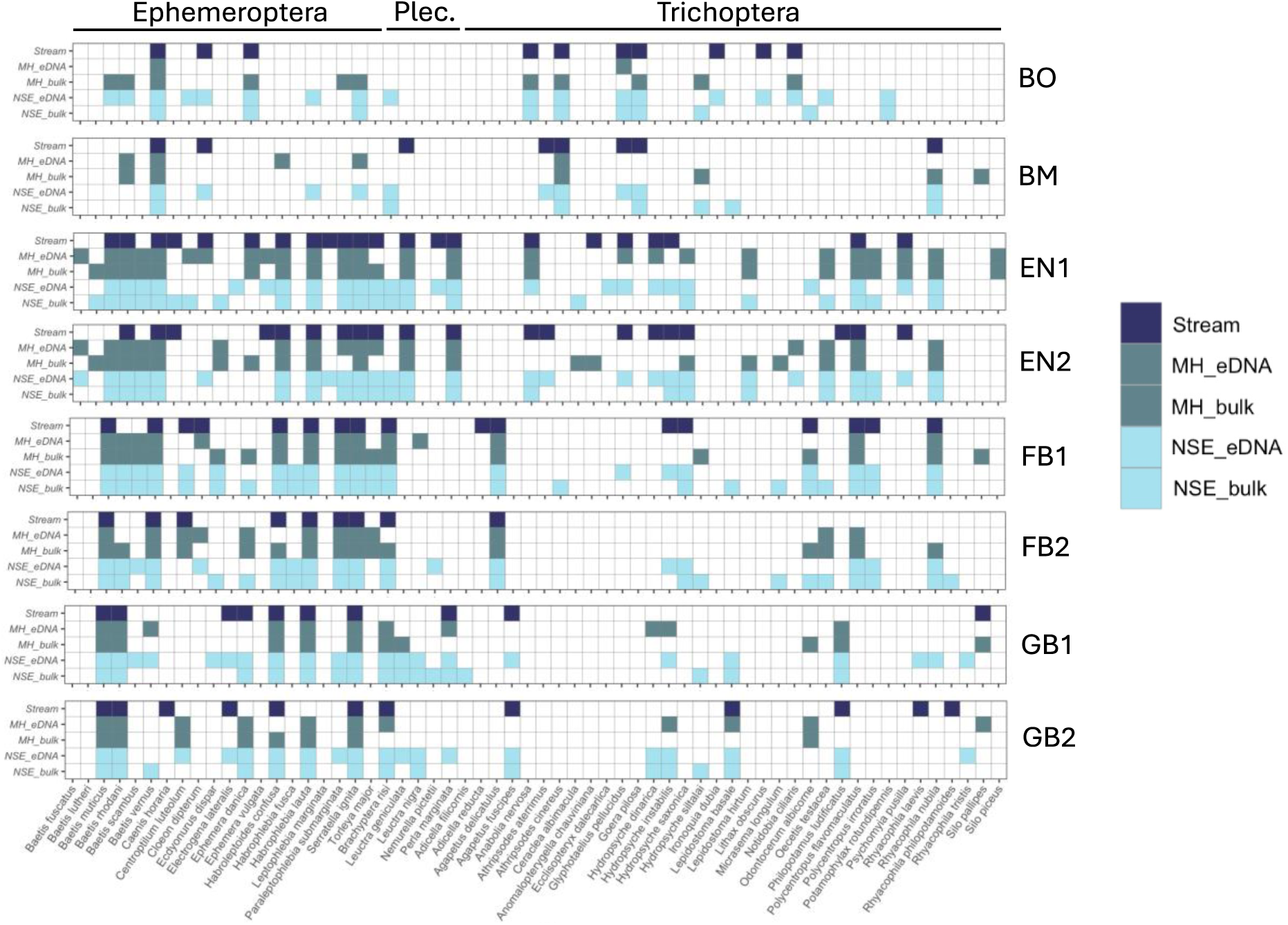
Detection of Ephemeroptera, Plecoptera and Trichoptera species with a record in the OTL for the stream water samples, multi-habitat kicknet (MH) eDNA and bulk samples and the natural substrate exposure (NSE) eDNA and bulk samples. Detection is displayed for each sampling site. Dark blue: Stream water eDNA samples, grey blue: MH eDNA and bulk samples, light blue: NSE eDNA and bulk samples.

## 4. Discussion

Our study demonstrates that metabarcoding of eDNA enriched water through incubation of locally sampled animals can reliably detect local macroinvertebrate stream communities, as typically done in the context of the Water Framework Directive (WFD). In particular, we showed that the use of preassembled natural substrate exposures (NSEs) colonised by the local invertebrate community were suitable to detect small-scale spatial differences in invertebrate trait composition between pool and riffle stream sections using both, metabarcoding of incubated water samples and bulk. Furthermore, ecological status class values obtained from metabarcoding of incubated eDNA samples, bulk samples and from stream water closely matched those values derived via independent morpho-taxonomic assessments. This highlights the general suitability of these approaches for ecological quality assessments. The only exception was eRNA metabarcoding, which consistently indicated poorer ecological status class values across sampling sites, indicating current limitations for its application for ecological quality assessment.

### 4.1 Does the natural substrate exposure (NSE) sampling approach provide an accurate reflection of invertebrate trait diversity in stream habitats?

While the multi-habitat (MH) sampling approach covered a broader range of sampled habitats and also a larger area (0.625 vs. 0.360 m^2^) than the natural substrate exposures (NSEs), the invertebrate trait composition did not differ significantly between the two approaches. As a result, the NSE sampling approach seems to reliably derive stream habitat diversity from the identified invertebrate community despite the lower diversity of covered habitats. The high similarity between sampling approaches found in this study, however, may also be due to the little variation in microhabitat and food resources at the sampled sites. Study sites were dominated by gravel substrates (60-85%), except for the Boye and thus differ from other stream types. The Boye was more diverse in its microhabitats with a large section (40%) of the streambed being covered in sand. These sandy substrates were not represented by the NSEs, which were limited to a wood-leaf mixture and gravel (Dumeier et al. 2018). We therefore expected a higher proportion of sand-associated species in the MH samples of the Boye, which we did not detect. *Ephemera danica* Müller, 1764, a typical sand-dwelling species, was detected throughout all streams, except for the Borbecker Mühlenbach. The stream is dominated by gravel substrates, similar to the other streams, and offers only a few sand patches. The absence of *E*. *danica* at the Borbecker Mühlenbach is likely caused by the “poor” ecological status of the site, as *E*. *danica* is known to be sensitive to stressors (Buffagni 1997). Sand is generally a habitat with lower species richness and abundance compared to gravel-dominated habitats, as it offers a less stable surface for attachment, biofilm growth, or shelter for non-sand-dwelling species (Duan et al. 2008, 2009; Mathers et al. 2023). Stream substrates in the Boye were dominated by sand but also included patches of gravel, macrophytes and wood. These habitats are known to offer a higher species richness and abundance than sand in sand-dominated lowland streams (Brusven and Prather 1974; Lorenz et al. 2004; Shupryt and Stelzer 2009; Mathers et al. 2024). This may explain the similar trait composition between the MH and NSE samples and the low species richness, especially for orders Ephemeroptera, Plecoptera and Trichoptera (EPT), in the Boye. Another possible explanation for the similar trait composition between sampling approaches is that the NSEs offer additional features that attract different species. For instance, the nets and their attachment to the streambed may provide a stable surface for attachment and protection from high discharge. Another factor to consider is the accumulation of suspended particulate matter on the surface of the NSEs containing DNA from species not present within the NSEs. However, we did not detect differences in read abundances between MH and NSE samples for sand-dwelling species.

When analysing the four NSEs per stream individually, no significant difference in trait composition was observed between the two NSE substrate types wood-leaf and gravel, neither for microhabitat preference nor feeding type. This could be explained by the proximity of similar food resources and microhabitats near both NSE types. Additionally, biofilm growth was observed on both types of NSEs, which may have attracted similarly grazing invertebrates to both NSE types. In contrast, trait composition differed significantly between NSEs placed at distinct flow regimes. Species preferring low flow currents were more prominent in the pool habitat samples, while those preferring high flow currents were predominantly found in the riffle habitat samples. These results align with the different environmental conditions found in pool and riffle habitats, which also influences the occurrence of different feeding types. In accordance with literature, we found a higher proportion of grazer species in riffle samples, likely due to increased biofilm growth supported by a high oxygen concentration and reduced settlement of sediments on the gravel surfaces (Cummins 2016). In contrast, we found a higher proportion of gatherer species at the pool samples, as sediment settles down in pools due to the reduced current (Cummins 2016). We detected more passive filter feeders in riffles, which rely on a high flow for filtering. Conversely, we detected more active filter feeders in pools, likely because they can filter independently of the flow current by actively creating currents around themselves (Merritt and Wallace 1981). The higher proportion of predators found in riffles is not as evident but is likely related to the traits of their prey species and that in general riffle habitats have a higher species richness and abundance than pool habitats (Scullion et al. 1982; Logan and Brooker 1983; Brown and Brussock 1991). One notable finding was the parasite *Epoicocladius ephemerae* which was more associated with pool samples but according to literature prefers stronger currents (Schmidt-Kloiber and Hering 2015, 2024). The current preference trait, however, was only assigned based on subfamily level, because species trait information was unavailable. Its host, *Ephemera danica*, prefers lower flows and was predominantly found in the pool samples, suggesting that the parasite follows the habitat preference of its host (Šulc and Zavřel 1924; Schmidt-Kloiber and Hering 2015, 2024). This was also reflected in the detection of *E. ephemerae* in 86% of the bulk samples that also found *E. danica*. For the incubated water eDNA samples it was only 33%. Overall, trait composition showed stronger flow regime differences based on bulk DNA samples than based on incubated water eDNA samples. This indicates that the eDNA signal from the stream water was still detected as a background signal in the incubated water eDNA samples. Another possible explanation is that some locally occurring taxa at the site were not detected via eDNA from the incubation medium. These results align with the results from the community composition analysis. However, the rheophilic genus *Rhyacophila* was predominantly found in riffle samples of the incubated water eDNA samples, highlighting that metabarcoding of eDNA enriched water from locally sampled animals can detect flow-associated taxa at small spatial scales. Detecting such small-scale spatial patterns from stream water samples directly is unlikely. High water movement in smaller streams lead to fast intermixing between stream habitats, resulting in rapid DNA transport and mixing (Shogren et al. 2017; Fremier et al. 2019). Small scale differences in fish community composition along the lateral and longitudinal scale of a stream has been detected using eDNA metabarcoding of water samples from wider streams (e.g., Berger et al. 2020; Thalinger et al. 2021). However, such local differentiation has not been achieved for macroinvertebrates with eDNA metabarcoding for smaller streams with various pool and riffle habitats like in this study. Our results suggest that NSEs can effectively capture stream habitat diversity by using both, bulk and incubated water eDNA metabarcoding, but performance of NSEs can vary depending on the placement within different flow regimes. For ecological status assessment using “Perlodes”, the number and placement of NSEs in stream sections with different flow conditions are probably more important for reflecting the actual stream habitat diversity than NSE substrate composition.

### 4.2 Are ecological status class (ESC) values derived from metabarcoding of eDNA enriched water through incubation of locally sampled animals consistent with morpho-taxonomic assessments?

Ecological status class (ESC) values derived from taxa lists obtained by metabarcoding of incubated water eDNA, bulk DNA and stream water eDNA were consistent between each other. Furthermore, they were consistent with routinely assessed morpho-taxonomic ESC values stored in the ELWAS database and with ESC values from the sampling campaign conducted by expert taxonomists at the site. Although more Ephemeroptera, Plecoptera and Trichoptera (EPT) species were detected using incubated water eDNA and bulk metabarcoding, the resulting ESC values did not differ by more than one class among metabarcoding approaches or in comparison to the morpho-taxonomic assessments. Metabarcoding can potentially result in the estimation of higher ecological status classifications through improved detection of smaller taxa (Múrria et al. 2024). However, in this study, the high congruence between presence/absence-based metabarcoding and the abundance-based morpho-taxonomic assessments demonstrates the robustness of metabarcoding for ESC assessments, also shown in the study by Macher et al. (2025). This is further supported by the incubated water eDNA sample from the MH approach of the Boye, which, despite detecting only six taxa, still assigned a “moderate” ESC value comparable to the other approaches. This indicates that even a limited number of indicator taxa can suffice to assess the ecological status of a stream. The low taxa richness in the incubated water eDNA samples from the Boye was likely caused by sediment inflow from a nearby construction, which impaired water filtration of the incubation medium. In contrast to the NSE sampling approach, the MH sampling approach disturbed the streambed, stirring up sediment that was captured in the sampling net. This sediment was then transferred to the incubation medium along with the sampled animals, likely reducing filter permeability.

Particularly in the Emscher tributaries (Boye and Borbecker Mühlenbach), deviations between ESC values derived from the taxa lists of the four NSEs of a sampling site were more pronounced than in the other stream sites. These discrepancies did not systematically differ based on NSE type or placement at different stream habitats, suggesting random variation. The differences may be the result of still ongoing recovery processes of the stream communities, as the Emscher catchment was only recently restored after serving as a wastewater channel. This is also supported by the still poor ecological status of the Borbecker Mühlenbach sampling site which has been wastewater free since 2014. The invertebrate communities at such sites are likely still in transition, with fluctuating species composition due to ongoing recolonization and adaptation as the recovery of freshwater communities takes on average between 10 and 20 years (Jones and Schmitz 2009; Winking et al. 2014; Gillmann et al. 2025). Given this dynamic state in recently restored streams, regular monitoring using minimally invasive approaches is crucial to monitor restoration success and to inform on management strategies. Therefore, the implementation of metabarcoding and especially eDNA-based metabarcoding into regulatory biomonitoring in the context of the WFD can act as an important monitoring tool for environmental decision making.

### 4.3 Potential of the different metabarcoding approaches for the implementation into the WFD

All presented approaches consistently assessed the ecological status class of the sampled stream sites with the exception of stream water eRNA metabarcoding. The persistence and degradation dynamics of eRNA are not yet well understood and range from similar persistence to eDNA to a much higher degradation rate (Mengoni et al. 2005; Wood et al. 2020; Marshall et al. 2021; Jo et al. 2023; Jo 2024). We observed a low taxa richness in many of our eRNA samples, which likely impaired ESC calculations. However, in some cases, for example for one sample site of the Grüner Bach, we detected the same ESC value as with the morpho-taxonomic assessment despite the limited taxa richness and only one available replicate. These findings highlight the need for more research regarding eRNA metabarcoding, particularly on its stability in the environment and processing before it can be considered for routine application into the WFD. Nonetheless, the analysis of eRNA holds considerable potential for future local impact assessment. Particularly for assessing the physiological status and health of individuals or their ecological stress response following a local impact (Ikert et al. 2021). Here, eRNA analysis has the potential to function as a predictive tool for the detection of changes at the molecular level, before those changes can be assessed via eDNA or traditional biomonitoring approaches.

Although we detected fewer EPT indicator taxa for ecological status class assessments of the stream water eDNA samples, the status class values were generally consistent with those from the independent morpho-taxonomic assessment. This indicates that eDNA metabarcoding of a direct stream sample has potential for future implementation under the WFD. The approach offers several advantages, including an easy and fast sampling and the ability to sample a broad spectrum of biodiversity across taxonomic groups, from microbes and invertebrates to fish and mammals and even terrestrial species (Deiner et al. 2016; Macher et al. 2021a; Altermatt et al. 2025). However, this holistic approach limits the potential of the approach to assess the local invertebrate community, as required by the WFD’s standardised, site-specific sampling protocols (see Figure 7 for an overview of the advantages and disadvantages of the approach). Currently, the new sampling procedures are not implemented into the WFD and can only be considered in a revision process, which will not happen before 2027 (Hering et al. 2018; Altermatt et al. 2025). However, the holistic approach and regional nature of the eDNA sampling approach could serve as a complementary tool in future monitoring frameworks. The restriction of the “Perlodes” approach to stream sections of a defined length (20-50 m for catchment sizes of 10-100 km^2^ and 50-100 m for catchment sizes of 100-10.000 km^2^) is mostly due to the high effort required for the monitoring of entire stream catchments using the standard sampling procedures (Meier et al. 2006). In the future, this limitation may be overcome by eDNA-based approaches, offering a broader spatial coverage with reduced sampling effort.

**Figure 7.**
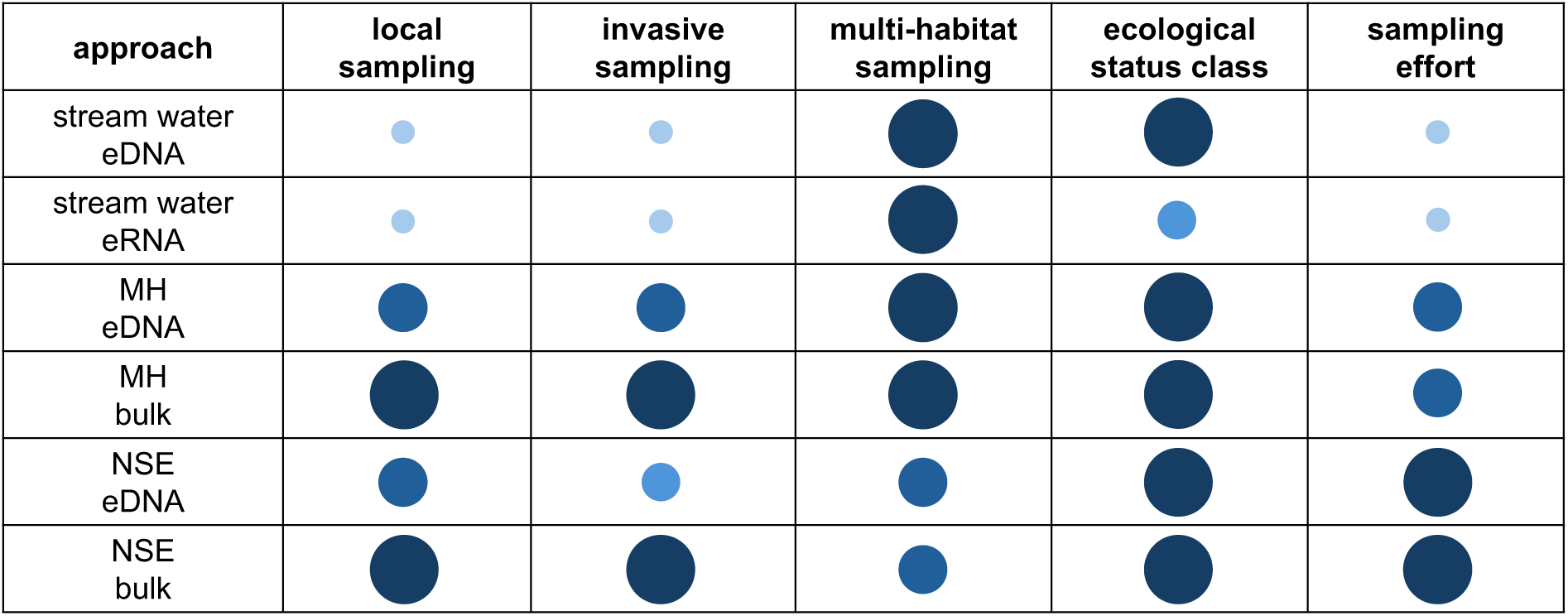
Rating of different criteria for monitoring of the macroinvertebrate stream community using the six sampling approaches analysed in this study. Circles indicate the weighting of the different criteria per sampling approach from large circles (high weight) to small circles (low weight).

For a more rapid implementation into the WFD and its current regulations, the incubation strategy appears particularly promising. The strategy is, similar to bulk metabarcoding, capable of detecting the local stream invertebrate community (Elbrecht et al. 2017; Meyer et al. 2021; Sander et al. 2025; Sander et al. in press). Bulk metabarcoding is already considered suitable for the implementation into the WFD in Germany using “Perlodes” without major adjustments (Macher et al. 2025). However, bulk metabarcoding involves the killing of the sampled animals for DNA extraction. In contrast, the incubation strategy offers a more animal-friendly alternative by allowing to release the collected animals back to the stream after incubation. This study demonstrated that both the incubation strategy and bulk metabarcoding detected similar invertebrate communities with a high similarity for taxa used in the “Perlodes” approach. Similarly, ESC values derived from both approaches were consistent with each other and with the morpho-taxonomic assessments. However, the multi-habitat (MH) sampling approach is still destructive to the streambed and stresses the sampled animals. To address this, the natural substrate exposure (NSE) sampling approach was proposed. The NSEs can be placed and retrieved without disturbing the streambed or the present animals and can be produced in a standardised format (De Pauw et al. 1986). A key limitation of the NSE approach is the inability to replicate certain stream habitats, like sand or other elusive substrates. While taxa richness and trait composition were not significantly different for the sand-dominant stream Boye for the NSE samples, species specialized in certain microhabitats or feeding types may be underrepresented in other stream types. Therefore, further research is needed to assess the applicability of the NSE approach for a broader range of habitats and stream types, as well as its potential as a standardised biomonitoring tool within the context of the WFD. One major advantage of the MH approach is that the sampling method used already aligns with the standardised sampling protocol used for the “Perlodes” approach and that it reflects the full range of a stream’s habitat diversity, facilitating its implementation into current frameworks compared to the NSE approach (see Figure 7 for a comparison of the advantages and disadvantages between the MH and NSE approach). In more simple terms it is a change in only a small step (i.e., the identification step) of the analysis workflow (Option 1 in Hering et al. 2018). Here, the MH approach combines a standardised sampling with animal welfare. Despite the possible limitations, the NSE approach holds particular value for recolonisation experiments. Dumeier et al. (2020) demonstrated the benefits of the NSEs for a successful and minimally invasive recolonisation of streams. The incubation strategy could complement these recolonisation experiments by analysing the transferred community of the donor stream using metabarcoding of the eDNA enriched water from the incubation medium before the NSEs are relocated. As a result, information on the donor invertebrate community can be obtained without compromising animal welfare, which is not possible with other approaches like morphological identification or tissue-based metabarcoding. The NSE approach may thus serve as a fast, minimally invasive tool for monitoring and thereby facilitating the recolonisation success of disturbed or recovering communities. In general, genetic methods should be considered, at minimum, as complementary tools for ecological status class assessment under the WFD. They offer important advantages, like a high taxonomic resolution, especially for taxonomically often neglected taxa like annelids and chironomids and can detect species- and OTU-specific responses to environmental stressors and seasonal variations (Resh and Unzicker 1975; Bista et al. 2017; Beermann et al. 2018; Sander et al. 2024). The incubation strategy in particular combines many of the advantages of bulk metabarcoding with the benefit of preserving animal welfare, reinforcing the importance for consideration for the implementation into the WFD.

## 5. Conclusion

Our study highlights the incubation strategy as a promising complementary tool for assessing the ecological status of local macroinvertebrate stream communities under the Water Framework Directive (WFD). The multi-habitat (MH) sampling approach of the incubation strategy, in particular, aligns well with the standardized procedures used in the “Perlodes” assessment. While further research is needed to evaluate and standardize natural substrate exposures (NSEs) across different stream types, the NSE sampling approach already shows potential for local impact assessments by detecting small-scale spatial differences in trait composition. Although direct stream water eDNA metabarcoding yielded comparable ecological status class values, its regional sampling design currently limits the integration into localized regulatory monitoring. Nevertheless, its broader sampling scope makes it a valuable complementary tool for future, catchment-scale assessments under the WFD. Overall, we encourage considering implementation of the incubation strategy as a complementary or alternative tool into the WFD as it offers a minimally invasive and localized sampling approach.

## Acknowledgements

We thank the Emschergenossenschaft und Lippeverband (EGLV) for their support for sampling the Emscher’s tributaries. We also thank Armin Lorenz for valuable input regarding construction and sampling of NSEs. MS was supported by the DBU scholarship program (20020/685). DB, FL, MW and AJB are members of and supported by the Collaborative Research Center (CRC) RESIST funded by the Deutsche Forschungsgemeinschaft (DFG, German Research Foundation) 1439/1— project number: 426547801.

## Data Availability Statement

All raw reads are accessible via the European Nucleotide Archive (ENA) under accession number PRJEB85525.

