## Appendix A for "Environmental DNA metabarcoding of invertebrate-incubated water supports WFD-compliant and animal-friendly bioassessment with added trait insights"

**Supplementary tables and figures**

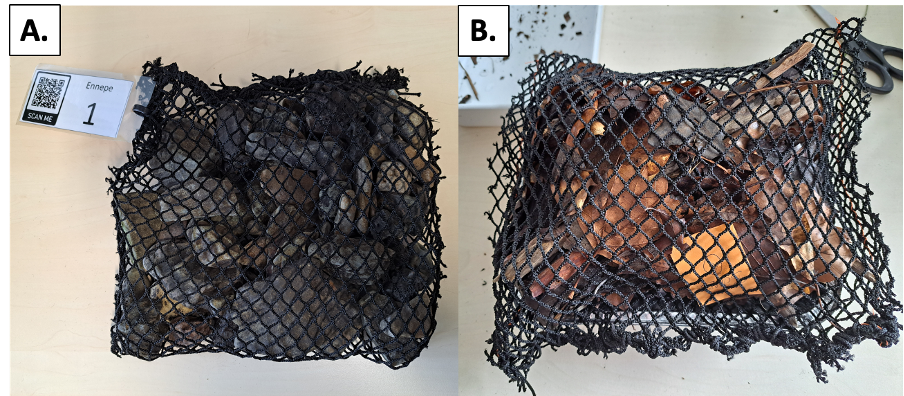

**Figure A.1.** Photo of the two NSE types before the attachment to the streambed. A: NSE filled with gravel, B: NSE filled with a mixture of wood and leaves. The substrates were held together with a nylon mesh.

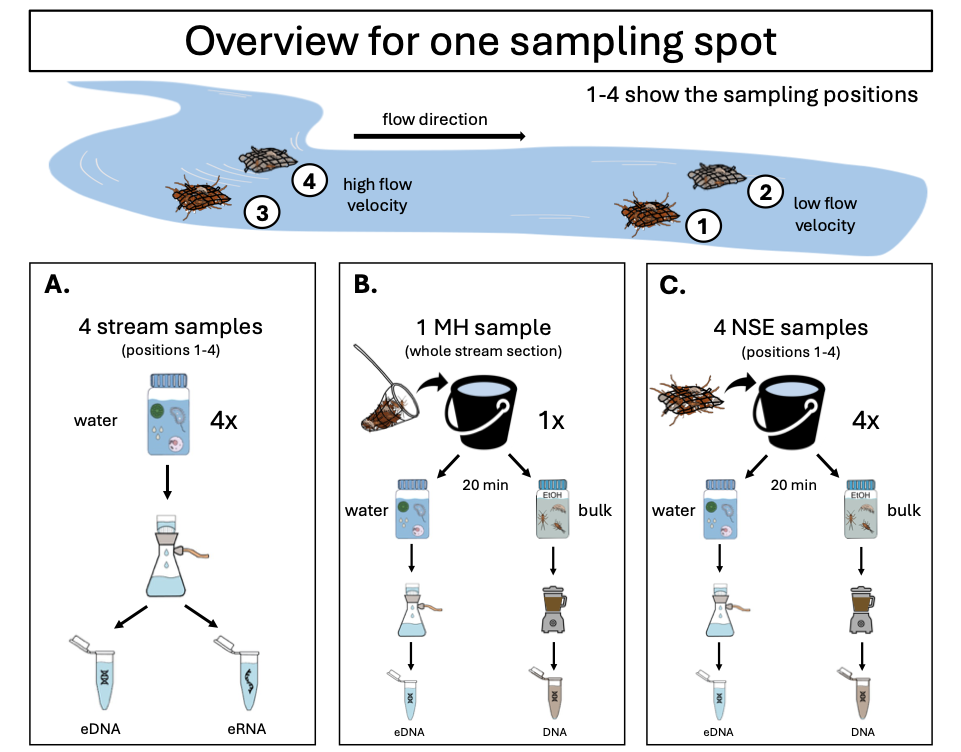

**Figure A.2.** Overview of the sampling, sample handling and DNA and RNA extraction approaches from the different samples for the three different sampling approaches: direct stream sampling (A); MH sampling (B) and NSE sampling (C). We sampled four streams: Emscher, Ennepe, Felderbach, Grüner Bach in Germany each at two sampling spots with four sampling positions for the stream water and NSE samples (32 sampling positions in total). For the MH samples, the whole stream section per sampling spot was sampled (8 stream sections in total reflecting the different microhabitats). A: stream water samples for eDNA/eRNA metabarcoding; B: incubation of MH samples and comparison of eDNA enriched in water to bulk MH DNA metabarcoding; C: incubation of NSEs and comparison of eDNA enriched in water to bulk NSE DNA metabarcoding. In total we analyzed 72 eDNA samples (32 stream eDNA, 32 NSE eDNA, 8 MH eDNA), 32 eRNA samples and 40 bulk DNA samples (32 NSE bulk, 8 MH bulk).

**Table A.1.** Read numbers for the different sample types after bioinformatic processing. Total number, mean number, minimum number and maximum number of reads. Note that for the MH samples only eight samples were used and for the stream and NSE samples 32.

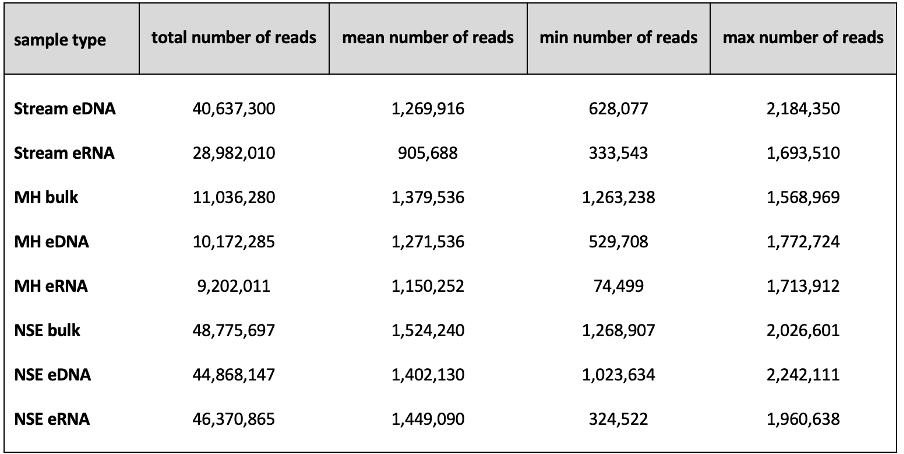

Note: We also analysed the enriched eRNA water samples from the incubation medium for both sampling approaches (MH and NSE). We did not use these data in this study but analysed them in the study by Sander et al. (in press).

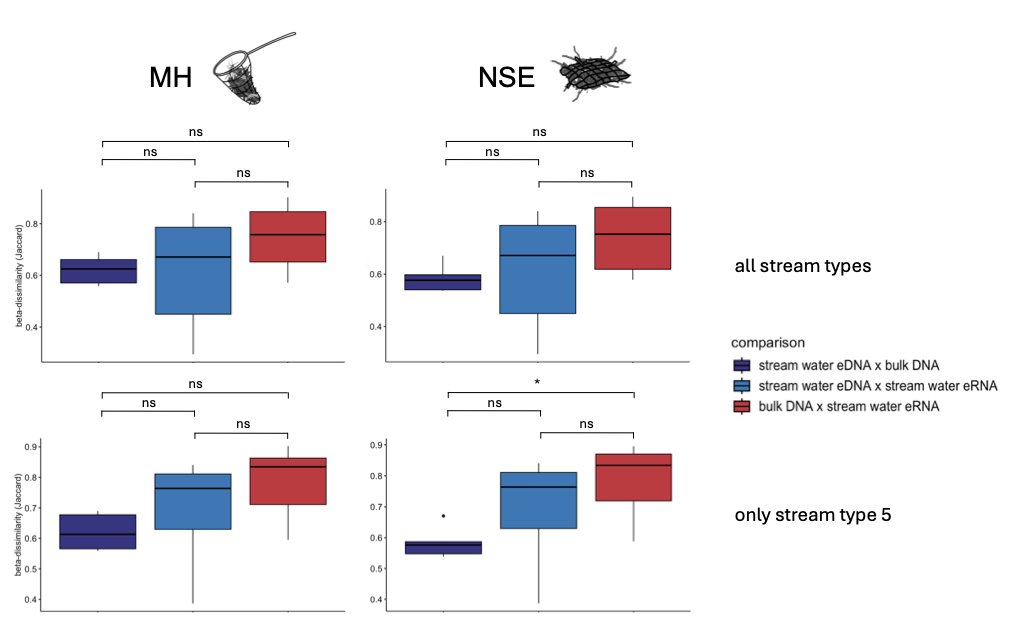

**Figure A.3.** Comparison of species community composition based on Jaccard dissimilarity between stream water eDNA, bulk DNA and stream water eRNA. Left: multi-habitat kicknet sampling approach (MH), right: natural substrate exposure sampling approach (NSE). Top: all streams, bottom: only streams assigned to stream type 5 according to LAWA typology. Significance between comparisons based on the results of the Tukey’s HSD test is presented as ns = not significant; * = p ≤ 0.05; ** = p ≤ 0.01; *** = p ≤ 0.001.

**Table A.2.** Indicator species for current preference for either pool or riffle NSE samples.

| **Species** | **Current preference** | **Habitat preference** | **A** | **B** | **Indicator value** | **p-value** |
| --- | --- | --- | --- | --- | --- | --- |
| ***Macrocyclops***  ***albidus*** | lip | pool | 0.90000 | 0.56250 | 0.712 | 0.0001*** |
| ***Acanthocyclops vernalis*** | lip | pool | 0.84211 | 0.50000 | 0.649 | 0.0009*** |
| ***Cypria  ophtalmica**** | lrp | pool | 0.81818 | 0.28125 | 0.480 | 0.0424* |
| ***Zavrelimyia  divisa**** | rlp | pool | 0.76190 | 0.50000 | 0.617 | 0.0079 ** |
| ***Oulimnius tuberculatus**** | rlp | pool | 0.80000 | 0.37500 | 0.548 | 0.0181 * |
| ***Dicranota  gracilipes**** | rlp | riffle | 0.81818 | 0.28125 | 0.480 | 0.0421 * |
| ***Dicranota***  ***pavida**** | rlp | riffle | 1 | 0.21875 | 0.468 | 0.0104 * |
| ***Epoicocladius ephemerae**** | rhp | pool | 0.90000 | 0.28125 | 0.503 | 0.0123* |
| ***Protonemura  praecox**** | rhp | riffle | 0.81818 | 0.56250 | 0.678 | 0.0003*** |
| ***Tvetenia***  ***verralli**** | rhp | riffle | 0.90909 | 0.31250 | 0.533 | 0.0067** |
| ***Agabus***  ***guttatus**** | rhp | riffle | 1 | 0.25000 | 0.500 | 0.0051** |
| ***Hydraena  gracilis**** | rhp | riffle | 0.76471 | 0.40625 | 0.557 | 0.0219* |
| ***Simulium  petricolum**** | rhp | riffle | 0.90000 | 0.28125 | 0.503 | 0.0139* |
| ***Simulium  variegatum**** | rhp | riffle | 1 | 0.21875 | 0.468 | 0.0109* |

Note: * = indicator species for pool or riffle for both current preference and feeding type. The indicator value is the product of the components A and B. A gives the probability of a site being a member of the site-group combination when the species has been found at that site. B informs of how frequently (and hence how easily) the species is found at sites of the site-group combination under study. Site is in this case the sample and group is the stream habitat (pool or riffle). Significance indicated as * = p ≤ 0.05; ** = p ≤ 0.01; *** = p ≤ 0.001.

**Table A.3.** Indicator species for feeding type for either pool or riffle NSE samples.

| **Species** | **Feeding type** | **Habitat preference** | **A** | **B** | **Indicator value** | **p-value** |
| --- | --- | --- | --- | --- | --- | --- |
| ***Oulimnius tuberculatus*†*** | gra/gat | pool | 0.80000 | 0.37500 | 0.548 | 0.0141* |
| ***Protonemura praecox*†*** | gra/gat | riffle | 0.81818 | 0.56250 | 0.678 | 0.0007*** |
| ***Orthocladius frigidus†*** | gra/gat | riffle | 0.75000 | 0.56250 | 0.650 | 0.0043** |
| ***Tvetenia***  ***verralli*†*** | gra/gat | riffle | 0.90909 | 0.31250 | 0.533 | 0.0055** |
| ***Hydraena***  ***gracilis**** | gra | riffle | 0.76471 | 0.40625 | 0.557 | 0.0222* |
| ***Lype***  ***phaeopa*** | gra | riffle | 0.90000 | 0.28125 | 0.503 | 0.0120* |
| ***Micropsectra junci†*** | gat/aff | pool | 0.72222 | 0.40625 | 0.542 | 0.0475* |
| ***Cypria***  ***ophtalmica**** | gat | pool | 0.81818 | 0.28125 | 0.480 | 0.0414* |
| ***Simulium petricolum**** | pff | riffle | 0.90000 | 0.28125 | 0.503 | 0.0148* |
| ***Simulium variegatum**** | pff | riffle | 1 | 0.21875 | 0.468 | 0.0104* |
| ***Zavrelimyia***  ***divisa**** | pre | pool | 0.76190 | 0.50000 | 0.617 | 0.0066** |
| ***Agabus***  ***guttatus**** | pre | riffle | 1 | 0.25000 | 0.500 | 0.0054** |
| ***Dicranota gracilipes**** | pre | riffle | 0.81818 | 0.28125 | 0.480 | 0.0408* |
| ***Dicranota***  ***pavida**** | pre | riffle | 1 | 0.21875 | 0.468 | 0.0109* |
| ***Epoicocladius ephemerae**** | par | pool | 0.9000 | 0.2812 | 0.503 | 0.0119* |

Note: * = indicator species for pool or riffle for both current preference and feeding type. † = indicator species for pool or riffle for two feeding type traits. The indicator value is the product of the components A and B. A gives the probability of a site being a member of the site-group combination when the species has been found at that site. B informs of how frequently (and hence how easily) the species is found at sites of the site-group combination under study. Site is in this case the sample and group is the stream habitat (pool or riffle). Site is in this case the sampling site and group the stream habitat pool or riffle. Significance indicated as * = p ≤ 0.05; ** = p ≤ 0.01; *** = p ≤ 0.001.

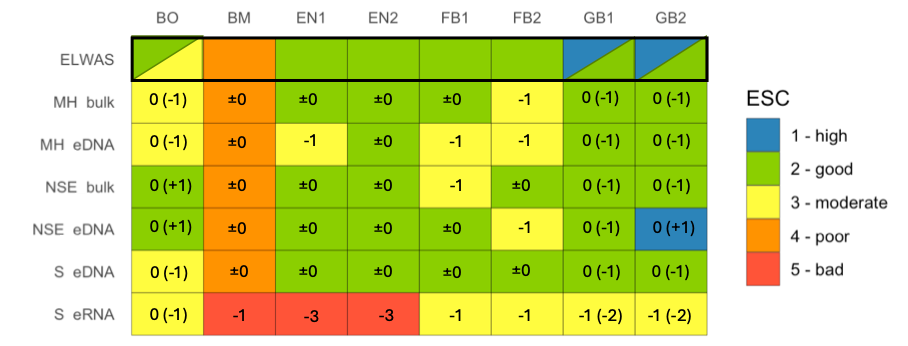

**Figure A.4.** Intercalibrated ecological status class (ESC) values calculated from taxa lists generated via metabarcoding and morphology-based taxa lists. Intercalibration was performed following the method used in Macher et al. 2025. ESC values range from 1 = high to 5 = bad. Values in brackets indicate the divergence of the value calculated via metabarcoding from the value stored in ELWAS. For sampling sites BO no recent morphology-based value was available at ELWAS. The values displayed are based on values extracted from ELWAS from the years 2012-2014. In addition, data from routine monitoring at this site reported an ESC value between 2 and 3. For the two sites GB1 and GB2, the most recent entry from ELWAS reports an ESC value of 1, however, no taxa list was available to validate the result. Therefore, we also included the older value stored with taxa lists in ELWAS, which report an ESC of 2.

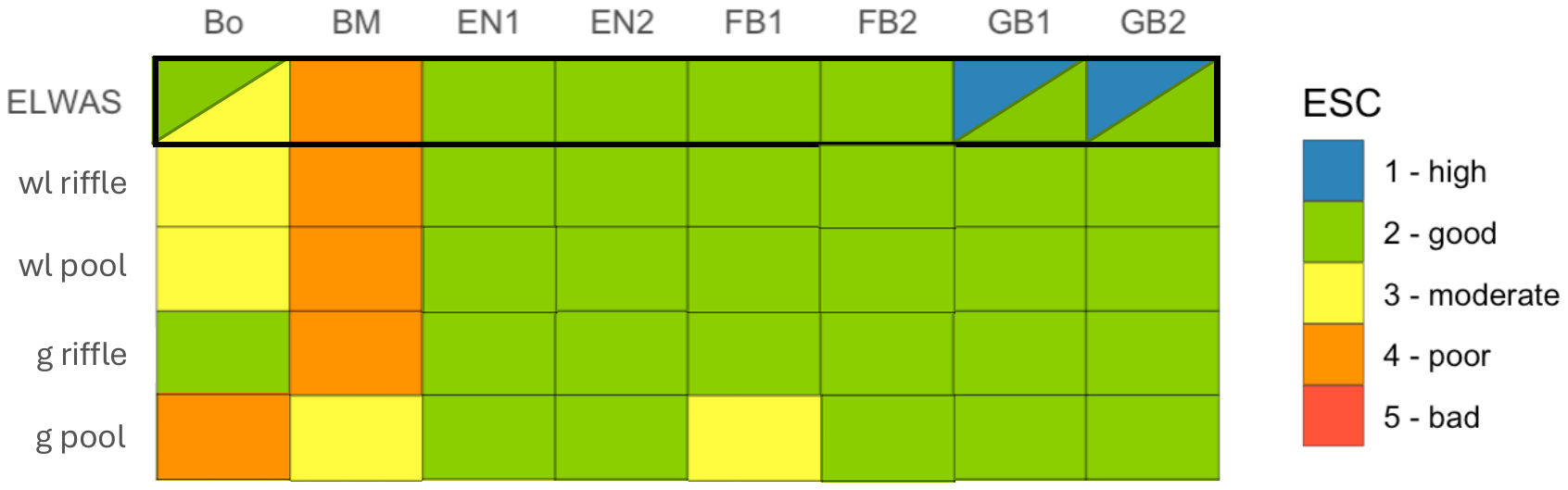

**Figure A.5.** Ecological status class values calculated from metabarcoding of NSE bulk DNA taxa lists. ESC values were calculated from each natural substrate exposure (NSE) separately. wl = wood-leaf NSE; g = gravel NSE; riffle = NSEs placed at riffle habitats; pool = NSEs placed at pool habitats. ESC values range from 1 = high to 5 = bad.

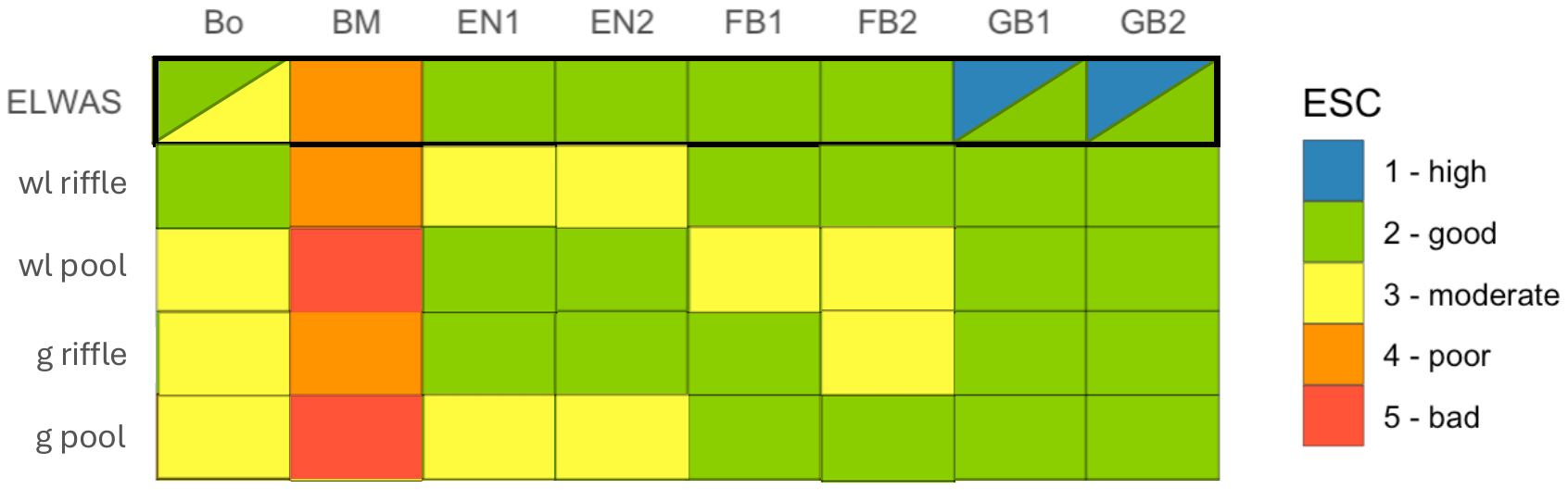

**Figure A.6.** Ecological status class values calculated from metabarcoding of NSE incubated water eDNA taxa lists. ESC values were calculated from each natural substrate exposure (NSE) separately. wl = wood-leaf NSE; g = gravel NSE; riffle = NSEs placed at riffle habitats; pool = NSEs placed at pool habitats. ESC values range from 1 = high to 5 = bad.
